# PGM3 inhibition rewires RUVBL2-dependent DNA repair and induces a BRCAness-like state in pancreatic cancer cells

**DOI:** 10.64898/2026.08.31.746486

**Authors:** Barbara Zerbato, Gabriele Taverna, Marina La Chimia, Marilena Pontoriero, Silvia Lombardi, Lorenzo Taglietti, Ke Deng, Giulia C.M. Perrone, Sini Hakkola, Alma Vuori, Teemu Syriälä, Emmanuel De Billy, Silvia Barabino, Cinzia Bragato, Ciro L. Pierri, Barbara La Ferla, Alfonso Urbanucci, Domenica Scumaci, Ferdinando Chiaradonna

## Abstract

Pancreatic ductal adenocarcinoma (PDAC) exhibits profound metabolic rewiring and strong resistance to DNA-damaging therapies, yet how metabolic pathways regulate genome maintenance remains poorly understood. The hexosamine biosynthetic pathway (HBP) integrates nutrient availability with protein glycosylation through production of UDP-GlcNAc, but its role in DNA damage response (DDR) regulation is unclear. Here we show that inhibition of the HBP enzyme phosphoglucomutase-3 (PGM3) reduces DNA repair capacity in pancreatic cancer cells. Transcriptomic and functional analyses reveal that the selective PGM3 inhibitor FR054 amplifies gemcitabine-induced replication stress, disrupts ATR-CHK1 and ATM-CHK2 checkpoint signaling, and selectively impairs homologous recombination. Glycoproteomic profiling identifies the AAA+ ATPase RUVBL2 as a key metabolic-DDR node. Gemcitabine increases RUVBL2 O-GlcNAcylation, with Thr81 identified as a modified residue within the Walker A nucleotide-binding motif. Structural modelling predicts that Thr81 O-GlcNAcylation stabilizes the RUVBL1-RUVBL2 complex without compromising ATP-Mg engagement. PGM3 inhibition and Thr81 mutation similarly reduced ATR and ATM abundance and promoted persistent DNA damage, supporting a role for RUVBL2 Thr81 O-GlcNAcylation in sustaining checkpoint signalling and genome stability. Consequently, PGM3 inhibition induces a BRCAness-like state that sensitizes pancreatic cancer cells to PARP inhibition, both in vitro and in vivo, as well as to ionizing radiation. These findings reveal a nutrient-sensitive mechanism linking protein glycosylation to genome maintenance and identify HBP-dependent DNA repair as a potentially actionable vulnerability in pancreatic cancer.

## Introduction

Pancreatic cancer (PC) remains one of the deadliest malignancies and is widely expected to become the second leading cause of cancer-related death by 2030 in the United States^1^. Most patients present with locally advanced or metastatic disease, and more than 80% are not candidates for curative resection. For advanced pancreatic ductal adenocarcinoma (PDAC), current first-line systemic treatment typically includes modified FOLFIRINOX, gemcitabine (GEM) plus nab-paclitaxel, or, in selected suitable patients, NALIRIFOX, whereas GEM-based regimens remain important options in less suitable patients and in specific clinical settings^2,3^. However, both intrinsic and acquired resistance substantially limit GEM efficacy^4^. Reported mechanisms include altered drug uptake and metabolism, activation of pro-survival signaling pathways, metabolic rewiring, and enhanced DNA damage response and repair, underscoring the need for strategies that hinder these adaptive defences^4^.

PC cells undergo profound metabolic reprogramming, including increased reliance on glucose and glutamine metabolism together with enhanced de novo lipogenesis^5,6^. A fraction of glucose- and glutamine-derived carbon and nitrogen is diverted into the HBP, which generates UDP-N-acetylglucosamine (UDP-GlcNAc), the essential donor substrate for N-glycosylation and O-GlcNAcylation^7^. Within this pathway, phosphoglucomutase 3 (PGM3) catalyses the interconversion of GlcNAc-6-phosphate and GlcNAc-1-phosphate, thereby contributing to the control of intracellular UDP-GlcNAc pools and downstream glycosylation reactions^8,9^. Increased HBP flux supports tumor growth, contributes to chemoresistance, and has been implicated in GEM resistance in pancreatic cancer^10,11^. As a nutrient-sensing pathway, the HBP links cellular metabolism to signaling, cell-cycle progression, stress adaptation and, more recently, regulation of the DNA damage response^12^. In this context, O-GlcNAcylation functionally intersects with phosphorylation to modulate the recruitment, amplitude and activity of DNA repair signaling networks^12^. Consistent with this, the enzyme that catalyses O-linked β-N-acetylglucosamine (O-GlcNAc) addition is recruited to sites of DNA damage, and disruption of O-GlcNAc cycling compromises DNA lesion resolution and may impair homology-directed repair^12–14^. Moreover, O-GlcNAcylation at damage foci restrains the spread of early DNA damage signaling, consistent with an antagonistic or an agonistic interplay with phosphorylation-dependent events during the double-strand breaks response^13^.

Targeting the HBP is therefore an attractive strategy in combination with chemotherapy, as it may limit adaptive stress responses that sustain tumor cell survival and reduce treatment efficacy. Consistent with this rationale, we developed a small-molecule inhibitor targeting PGM3 and previously showed that both pharmacological and genetic restriction of HBP flux perturb protein glycosylation and induce cancer cell death^11,15,16^. These observations suggest that HBP inhibition disrupts not only metabolic homeostasis, but also stress-response networks that may be required to resist genotoxic challenge^16,17^. Given the emerging link between HBP activity, O-GlcNAc-dependent signaling and DNA damage repair, we hypothesized that pharmacological targeting of this pathway could sensitize pancreatic cancer cells to GEM by weakening adaptive stress responses and impairing the resolution of treatment-induced DNA damage.

Here, we identify a functional link between HBP activity and DNA repair in pancreatic cancer cells. We show that the selective PGM3 inhibitor FR054 exerts marked single-agent antitumor activity and synergizes with GEM across multiple pancreatic cancer models. FR054 suppresses homologous recombination (HR) and disrupts the ATR-CHK1 replication-stress checkpoint, leading to persistent and unresolved replication stress. Mechanistically, these effects are associated with altered O-GlcNAcylation and nucleocytoplasmic distribution of RUVBL2, an AAA+ ATPase required for PIKK-complex homeostasis^18,19^. By inducing a BRCAness-like state, FR054 also sensitizes pancreatic cancer cells to olaparib in vitro and in vivo and enhances their response to ionizing radiation. Together, our findings establish HBP inhibition as a strategy to disrupt DNA damage response (DDR) signaling and counteract therapy resistance in pancreatic cancer.

## Results

### HBP inhibition counteracts GEM-induced adaptive programmes and activates DNA damage responses

To define the effects of HBP inhibition and GEM exposure on PC cells transcriptomes, we performed RNA-seq in MIAPaCa-2 and BxPC3 cell lines treated with FR054, GEM, or their combination (Fig. 1a, b). Both models exhibited extensive transcriptional reprogramming, with the combination inducing the highest number of differentially expressed genes (DEGs log_₂_FC ≥ 2, FDR < 0.05). In MIAPaCa-2 cells, FR054-treated conditions (alone and in combination) showed a substantial overlap of DEGs compared to GEM alone (Fig. 1c), indicating a dominant transcriptional contribution of HBP inhibition. A similar trend was observed in BxPC3 cells (Fig. 1d); however, in this model, DEGs induced by the combination showed limited overlap with those induced by either single treatment, suggesting a more distinct and less integrated transcriptional response. These differences point to cell line-specific dependencies in response to treatment. In fact, hierarchical clustering of high-confidence DEGs (log_₂_FC ≥ 2, FDR ≤ 10^-5^, MIAPaCa-2: 1488 genes; BxPC3: 1026 genes; Supplementary Table 1 and 2) revealed clear separation of treatment groups, highlighting pronounced transcriptional divergence across conditions (Fig. 1e). This difference was further substantiated by comparing of DEGs across both cell lines and treatment conditions, which revealed a limited shared core of 340 genes (Supplementary Tables 3). Notably, the majority of these (275 genes, Supplementary Tables 4) were specific to the COMBO condition, indicating that only the combined treatment drives a convergent transcriptional program across cell models, distinct from single-agent effects (Extended Data Fig. 1a, b).

**Fig. 1:**
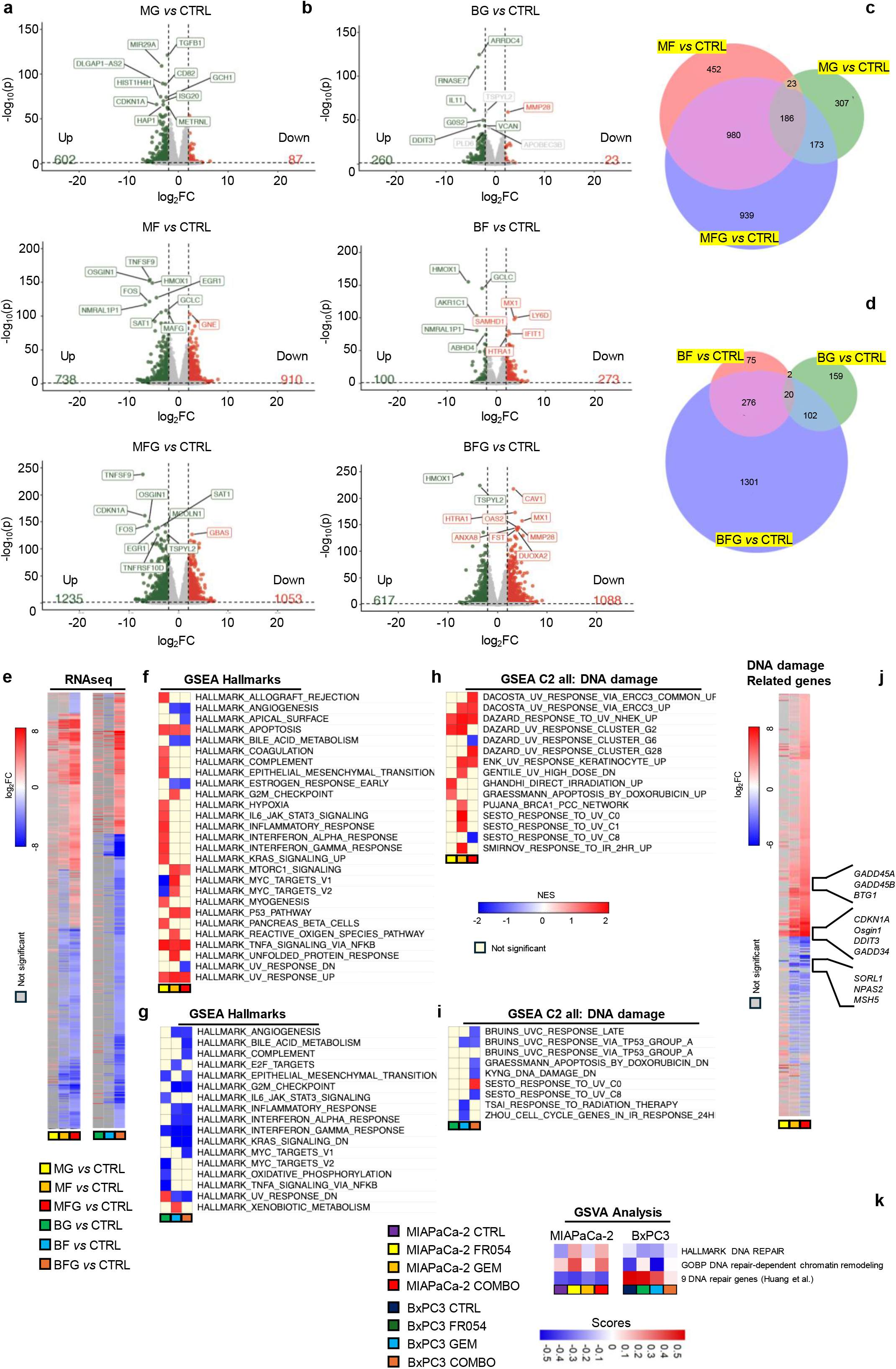
Transcriptomic and Gene Set Enrichment Analysis of MIAPaCa-2 and BxPC3 pancreatic cancer cells shows significant differences between the two cell lines upon FR054, GEM and COMBO treatments. **a, b,** Volcano plots of MIAPaCa-2 and BxPC3 pancreatic cancer cells treated for 48 h with 500 μM FR054, 1μM gemcitabine (GEM), and their combination (COMBO). The log_2_FC indicates the mean expression level for each gene. Each dot represents one gene. Gray dots, no significant DEGs between treated groups and control (CTRL) group, the green dots represent down-regulated genes, and red dots represent up-regulated gene, DEGs (log_2_FC>2, FDR<0.05). **c, d**, Venn diagrams illustrating the number of differentially expressed transcripts for different pairwise comparisons in MIAPaCa-2 and BxPC3 pancreatic cancer cells. **e**, Clustering of the DEGs identified in MIAPaCa-2, 1489 genes, and BxPC3, 1026 genes, showing a log_2_FC>2 and a FDR<10^-5^. Gray color, no significant DEGs between treated groups and CTRL group, the blue represents down-regulated genes and red represents up-regulated genes. **f, g**, Gene Set Enrichment Analysis (GSEA) performed by using the Hallmark gene sets and DEGs identified (log_2_FC>2, FDR<0.05) in MIAPaCa-2 and BxPC3 pancreatic cancer cells. Each box represents a Hallmark pathway in the three different conditions, as described in e, ranked by the Normalized Enrichment Score (NES). Positive NES values (red) indicate enrichment in the upregulated condition, while negative NES values (blue) indicate enrichment in the downregulated condition. The significance of enrichment was determined by FDR with pathways considered significant at FDR< 0.05. In light yellow not enriched pathway in the analyzed condition. **h, i**, GSEA performed by using C2 all and capturing only the pathways related to DNA damage inducing conditions by using DEGs identified (log_2_FC>2, FDR<0.05) in MIAPaCa-2 and BxPC3 pancreatic cancer cells. **j**, Clustering of 320 DEGs (log_2_FC>2 and FDR< 0.05) identified in MIAPaCa-2 GSEA C2 all: DNA damage. Gray color, no significant DEGs between treated groups and CTRL group, the blue represents downregulated genes and red represents up-regulated genes. Columns = samples. Rows = genes. Samples were clustered using Euclidean distance. Select genes associated with cell death and DNA damage labeled (right). **k**, GSVA heatmap of some DNA repair related processes inversely modulated in MIAPaCa-2 cells upon FR054 treatment as compared to BxPC3 cells.

To capture the full spectrum of transcriptional changes across both models, we performed Gene Set Enrichment Analysis (GSEA) using Hallmark gene sets^20^ on all high-confidence DEGs. In MIAPaCa-2 cells, GEM treatment enriched pathways associated with adaptive resistance, including epithelial-to-mesenchymal transition (EMT), KRAS signaling, and hypoxia (Fig. 1f). In contrast, FR054 treatment promoted enrichment of stress-related pathways, including p53 signaling, apoptosis, reactive oxygen species (ROS), unfolded protein response (UPR), and UV response. Notably, the combined treatment attenuated GEM-associated adaptive pathways, indicating that FR054 counteracts GEM-driven transcriptional adaptation (Fig. 1f). In BxPC3 cells, GSEA revealed a distinct profile characterized by a global reduction in pathway enrichment, particularly under combined treatment, consistent with a wide attenuation of transcriptional activity and potentially reflecting the greater sensitivity of this model to both GEM and the combined treatment (Fig. 1g).

Considering the prominent enrichment of stress-related pathways, especially in MIAPaCa-2 cells, and the well-established role of GEM as a DNA-damaging agent, we next focused on DDR using curated gene sets from the C2 collection^21^. In MIAPaCa-2 cells, both FR054 and combined treatments significantly enriched signatures associated with UV, ionizing radiation (IR), and double-strand break (DSB) responses, whereas these pathways were not enriched in BxPC3 cells (Fig. 1h, i). Clustering of 320 DNA damage-related DEGs in MIAPaCa-2 revealed robust induction of genes involved in negative cell proliferation control and induction of apoptosis, including GADD45A/B, BTG1, ATF3, DDIT3 and GADD34^22–26^ alongside downregulation of genes linked to chemoresistance, such as NPAS2, SORL1 and MSH5^27–29^ (Fig. 1j and Supplementary Table 5). To further detail our transcriptional analysis, we also performed Gene Set Variation Analysis (GSVA)^30^. This analysis again revealed divergent transcriptional arrangements between the two models: MIAPaCa-2 cells displayed coordinated activation of stress-associated programs under combined treatment, whereas BxPC3 cells exhibited a global reduction in pathway activity (Fig. 1k). Together, these findings indicate that combined treatment induces a stress-driven transcriptional reprogramming especially in MIAPaCa-2 cells, while promoting a more generalized transcriptional suppression in BxPC3 cells.

### HBP inhibition enhances GEM-induced cytotoxicity and DNA damage in pancreatic cancer cells

We previously demonstrated that pancreatic cancer cells rely on HBP activity to sustain proliferation under both metabolic^31^ and genotoxic stress conditions^11^, largely through enhanced protein N-glycosylation and activation of pro-survival signaling pathways. Consistently, inhibition of HBP disrupts these protective mechanisms, increasing cellular vulnerability to nutrient deprivation and to DNA-damaging agents such as GEM. Building on this rationale, and supported by the transcriptional changes observed above, we investigated whether the selective HBP inhibitor FR054 could enhance GEM efficacy by increasing cell death, impairing cell-cycle control and DNA damage resolution in pancreatic cancer cells characterized by K-ras mutation, namely MIAPaCa-2 and PANC-1 cell lines. Treatment of pancreatic cancer cells with FR054, GEM, or their combination resulted in marked cytotoxic and cytostatic effects. Indeed, trypan blue assays showed a time-dependent increase in cell death in MIAPaCa-2 (Fig. 2a) and PANC-1 cells (Fig. 2b), with the combination producing the strongest reduction, especially at later time point of analysis (72 h). Consistently, both FR054 and GEM impaired proliferation, while the combination maximally suppressed cell growth (Extended Data Fig 2a, c). Notably, FR054-treated cells appeared multinucleated, suggesting severe cytokinetic defects (Extended Data Fig 2b, d, yellow arrows).

**Figure 2.**
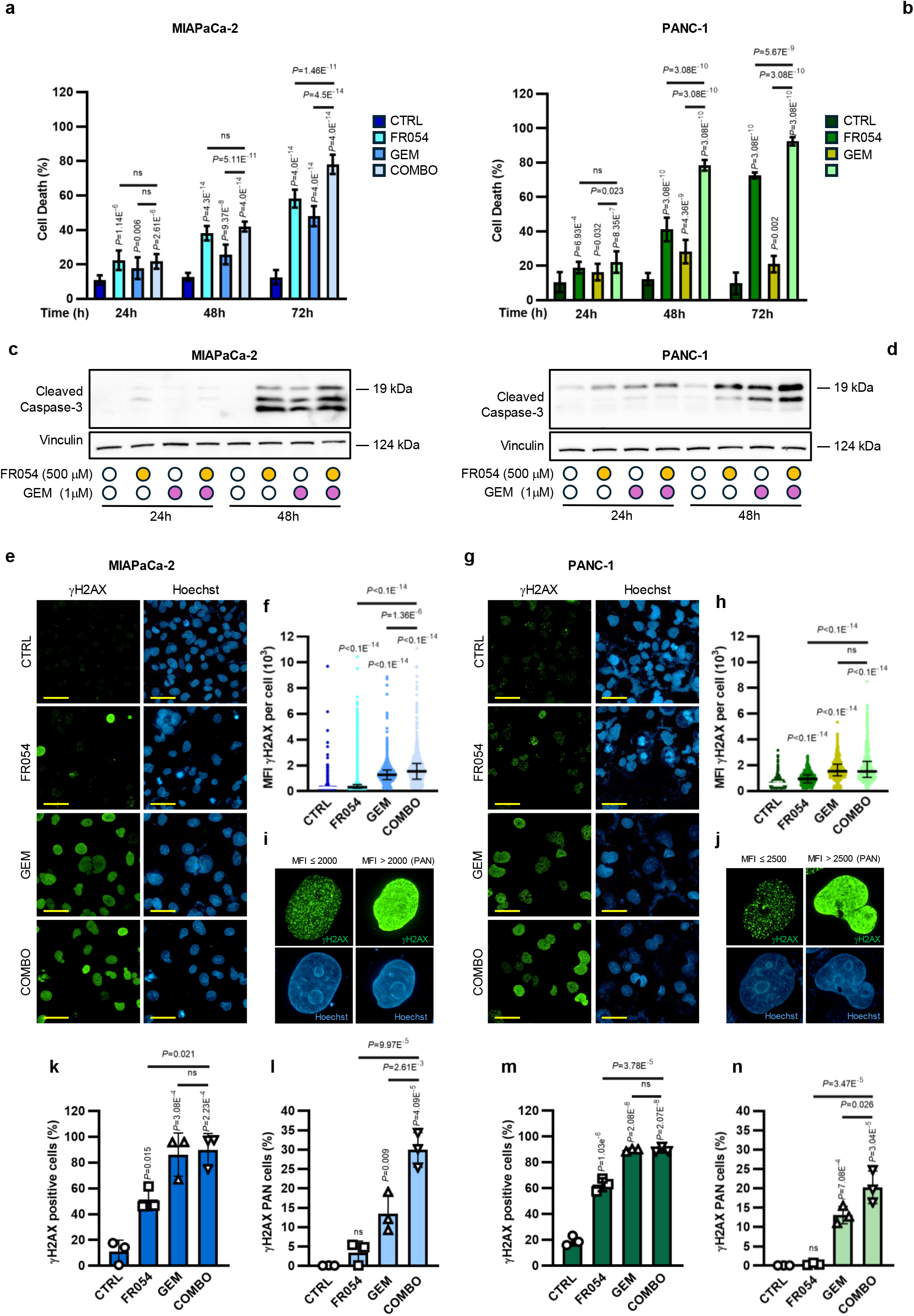
FR054 alone or in combination with GEM increases apoptosis in pancreatic cancer cells in association with an increased DNA damage. **a**, **b** Trypan blue vital assay in MIAPaCa-2 (**a**) and PANC-1 (**b**) cells treated for 24 h, 48 h and 72 h with vehicle (CTRL), 500 μM FR054, 1 μM gemcitabine (GEM), or their combination (COMBO). The data represents *n*=7–12 (**a**) and *n*=5–14 (**b**) biological replicates. **c**, **d**, Western blot analysis of cleaved caspase-3 in MIAPaCa-2 (**c**) and PANC-1 (**d**) cells after 24 h and 48 h of treatment with CTRL, FR054, GEM, or COMBO. Vinculin was used as a loading control. Representative immunoblots from three independent biological experiments are shown (*n*=3). **e, g**, Representative immunofluorescence images of γH2AX (green) and Hoechst (blue) at 48 h in MIAPaCa-2 (**e**) and PANC-1(**g**) cells treated as previously described. Scale bar, 50 μm (63× magnification). **f**, **h**, Single-cell quantification of γH2AX mean fluorescence intensity (MFI) in MIAPaCa-2 (**f**) and PANC-1 (**h**) cells using Harmony software after 48 h of treatments as described in **a**–**d**. *n*=3008–12530 (**f**) and *n*=1399– 4085 (**h**) cells were analysed per condition from *n*=3 biological replicates. **i**, **j**, Representative images of γH2AX-positive and PAN-positive cell populations identified according to MFI thresholds in MIAPaCa-2 (**i**) and PANC-1 (**j**) cells. PAN-positive cells were defined as cells with MFI > 2000 (**i**) or MFI > 2500 (**j**). **k**, **m**, Percentage of MIAPaCa-2 (**k**) and PANC-1 (**m**) γH2AX- positive cells treated for 48 h as described in **a**–**d** (*n*=3 biological replicates). **l**, **n**, Percentage of MIAPaCa-2 (**l**) and PANC-1 (**n**) PAN-positive cells treated for 48 h as in **a**–**h**, **k** and **m** (*n*=3 biological replicates). Data are presented as mean ± s.d. (**a**, **b**, **k**–**n**) or as scatter dot plots showing individual values, with the median indicated by the central line and the interquartile range (IQR) represented by the error bars (**f** and **h**). Statistical significance was determined using two-way ANOVA with Tukey’s multiple comparisons test (**a** and **b**), Kruskal-Wallis test with Dunn’s multiple comparisons test (**f** and **h**) or one-way ANOVA with Tukey’s multiple comparisons test (**k**–**n**). Exact *P* values are reported in the plots; ns, not significant.

Western blot analysis further revealed that reduced viability was accompanied by increased apoptosis, as shown by the induction of cleaved caspase-3 (Figure 2c, d).

We next assessed γH2AX staining by immunofluorescence at 24 h and 48 h. At both time points, all treatments markedly increased the mean γH2AX fluorescence intensity per cell and the percentage of γH2AX-positive cells compared with control conditions in both MIAPaCa-2 (Fig. 2e, f, k and Extended Data Fig. 2e, f, i) and PANC-1 cells (Fig. 2g, h, m and Extended Data Fig. 2g, h, j).

Notably, GEM and the combined treatment induced a significant increase in pan-nuclear γH2AX staining, a pattern associated with widespread chromatin damage. In MIAPaCa-2 cells, pan-nuclear staining was detected in approximately 13% and 20% of cells after GEM and combination treatment, respectively, at 24 h (Fig. 2i and Extended Data Fig. 2k), and in 12% and 30% at 48 h (Fig. 2i, l). In PANC-1 cells, the corresponding values were 25% and 23% at 24 h (Fig. 2j and Extended Data Fig. 2l) and 13% and 20% at 48 h (Fig. 2j, n). These findings indicate that the combination as compared to GEM alone, induces a time-dependent increase in widespread chromatin damage in MIAPaCa-2 cells and a more persistent response in PANC-1 cells.

Importantly, FR054 alone also increased γH2AX staining, supporting a role for the HBP in maintaining genomic stability. Overall, these results show that both FR054 and GEM promote DNA damage accumulation in pancreatic cancer cells, with the most pronounced effect observed following combined treatment.

### FR054 suppresses GEM-dependent ATR/ATM checkpoint activation and selectively impairs HR repair, leading to unresolved DNA damage

To elucidate the molecular mechanisms underlying the ability of FR054 to enhance GEM cytotoxicity, we next examined DDR pathways activated under single and combined treatments, given the strong accumulation of γH2AX alterations observed earlier. Consistent with previous studies^32,33^ in pancreatic cancer cells, GEM induced the expected replication-stress response, with time-dependent activation of the ATR-CHK1 axis, including phosphorylation of ATR (Thr1989) and Chk1(Ser345) (Fig. 3a, c), together with early RPA(Ser8) hyperphosphorylation (Fig. 3e, f). RPA phosphorylation declined by 48 h, indicating partial resolution of replication stress. Conversely, the combined treatment almost completely abrogated ATR and Chk1 activation in both MIAPaCa-2 and PANC-1 cells (Fig. 3a, c), while RPA(Ser8) persisted (Fig. 3e, f), demonstrating unresolved replication stress in the absence of functional checkpoint signaling. Of note FR054 alone did not elicit detectable ATR, CHK1 or RPA phosphorylation under these conditions, suggesting that the damage induced by HBP inhibition differs from the canonical replication-stress response triggered by GEM. Strikingly, FR054 treatment, alone or in combination, led to a marked reduction in total ATR protein levels as compared to GEM further supporting the checkpoint failure (Fig. 3a, c). Because ATR suppression after GEM exposure can convert stalled forks into DSBs and trigger compensatory activation of ATM, we analyzed ATM pathway engagement. GEM alone, especially at 24 h, robustly activated ATM(Ser1981) and its downstream target Chk2(Thr68) (Fig. 3b, d). In sharp contrast, FR054 in combination with GEM, strongly inhibited ATM phosphorylation, accompanied by a near-complete loss of total ATM protein (Fig. 3b, d). Interestingly, despite the marked reduction in ATM levels, Chk2 phosphorylation at Thr68 remained detectable in FR054-treated cells, although at a lower level than in GEM-treated cells (Fig. 3b, d).

**Figure 3.**
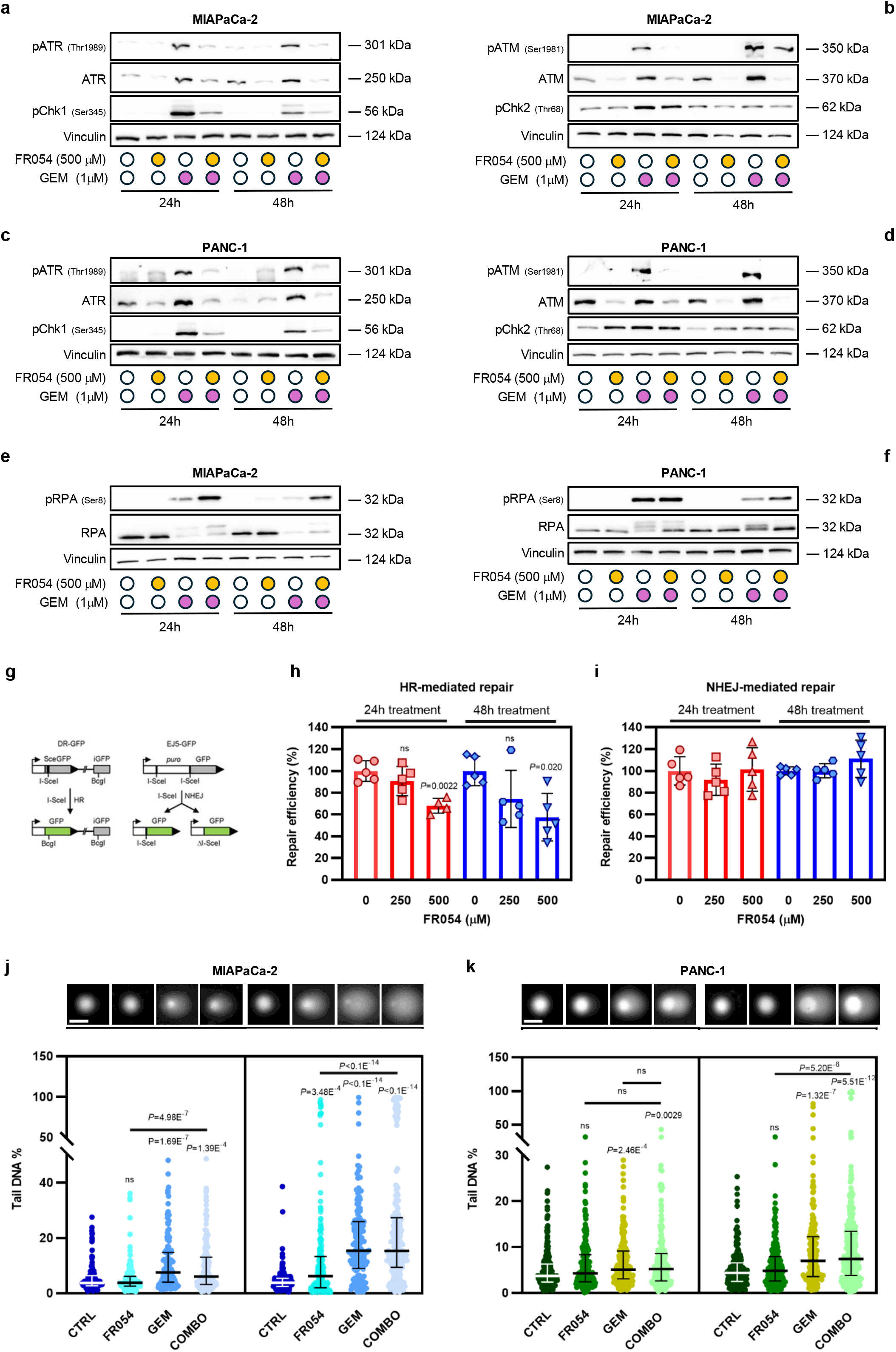
FR054 impairs ATR- and ATM-dependent DNA damage checkpoints, leading to defective HR and increased DNA damage. **a**, **c**, ATR pathway activation in MIAPaCa-2 (**a**) and PANC-1 (**c**) cells, respectively, treated with vehicle (CTRL), 500 μM FR054, 1 μM gemcitabine (GEM), or their combination (COMBO) for the indicated time points (24 h, 48 h). Protein levels of pATR (Thr1989), total ATR, and pChk1 (Ser345) were assessed by immunoblotting and vinculin was used as a loading control. Representative immunoblots from three independent biological experiments are shown (*n*=3). **b**, **d** ATM pathway activation in MIAPaCa-2 (**b**) and PANC-1 (**d**) cells, treated for 24 h and 48 h as previously described. Immunoblotting was performed to assess pATM (Ser1981), total ATM, and pChk2 (Thr68) levels, and vinculin was used as a loading control. Representative data from three independent experiments are shown (*n*=3). **e**, **f**, Immunoblot analysis of RPA hyperphosphorylation and pRPA (Ser8) levels in MIAPaCa-2 (**e**) and PANC-1 (**f**) upon treatments described in **a**, **c.** Representative blots from three independent biological experiments are shown (*n*=3) using vinculin as a loading control. **g**–**i**, HR (**h**) and NHEJ (**i**) repair efficiency analysis in U2OS reporter cells transfected with I- SceI-T2A-BFP plasmids (**g**) and treated with increasing concentrations of FR054 (250 μM and 500 μM) (**h**, **i**). HR (**h**) and NHEJ (**i**) repair activities were quantified from at least four independent biological replicates per condition (*n*≥4). FR054 treatment did not significantly affect NHEJ-mediated repair efficiency under the tested conditions. **j**, **k,** DNA damage analysis by comet assay in MIAPaCa-2 (**j**) and PANC-1 (**k**) cells after 24 h and 48 h of treatments as previously described in **a**, **c**. A total of *n*=130–201 (**j**) and *n*=188–295 (**k**) cells were analysed per condition from *n*=3 biological replicates. The data are presented as mean ± s.d. (**h** and **i**) or as scatter dot plots displaying individual values; the median is indicated by the central line and the interquartile range (IQR) by the corresponding bars (**j** and **k**). Statistical significance was determined using one-way ANOVA with Tukey’s multiple comparisons test (**h** and **i**) or Kruskal-Wallis test followed by Dunn’s post-hoc test (**j** and **k**). Exact *P* values are reported in the plots; ns, not significant.

We next investigated whether the suppression of ATR/ATM signaling was associated with defective cell-cycle progression. EdU incorporation was progressively reduced by FR054 and was almost completely abolished by GEM and the combined treatment at both 24 h and 48 h in MIAPaCa-2 and PANC-1 cells, indicating a stable inhibition of DNA synthesis (Extended Data Fig. 3a-d). Consistently, immunoblotting revealed distinct treatment-dependent changes in cyclins controlling successive phases of the cell cycle. FR054 alone mainly reduced Cyclin D1, consistent with a general antiproliferative effect. GEM decreased Cyclin D1 while increasing Cyclin E, Cyclin A and Cyclin B1, suggesting progression beyond G1 followed by accumulation in S and G2 in response to replication stress. The combined treatment maintained Cyclin D1 suppression but, especially at 48 h, significantly reduced Cyclin A and Cyclin B1 expression (Extended Data Fig. 3 e, f), suggesting a reduction in accumulation of cells in S and G2. To determine how the alterations in DNA synthesis and cyclin expression were related to DNA damage accumulation, we next performed a single-cell analysis of Cyclin A and γH2AX, a combination previously used to identify replication-associated damage in Cyclin A-positive S/G2 cells^34,35^. Single-cell population analysis revealed a marked treatment-dependent redistribution of cells according to Cyclin A and γH2AX levels (Extended Data Fig. 3g-j and Supplementary Fig. 1a, b). FR054 alone expanded the Cyclin A-high/γH2AX-high population, particularly in MIAPaCa-2 cells, indicating that HBP inhibition promotes DNA damage in proliferating cells. GEM shifted most cells towards a Cyclin A-high/γH2AX-high state, consistent with the accumulation of damaged cells in S-phase. The combined treatment produced distinct time-dependent responses in the two cell lines. In MIAPaCa-2 cells the Cyclin A-high/γH2AX-high population increased further at 48 h, suggesting persistence of damaged cells in an aberrant S-phase state (Extended Data Fig. 3g, h and Supplementary Fig. 1a). Conversely, in PANC-1 cells, the combination increased the Cyclin A-low/γH2AX-high population, indicating that DNA damage persisted despite the progressive loss of Cyclin A (Extended Data Fig. 3i, j and Supplementary Fig. 1b). Thus, FR054 prevents the normal resolution of GEM-induced damage, either by retaining cells in a damaged Cyclin A-positive state or by uncoupling persistent DNA damage from the maintenance of the S-phase programme. These findings, together with the marked reduction in EdU incorporation and checkpoint inhibition, indicate that FR054 interferes with the orderly progression of GEM-treated cells through S and G2/M, preventing completion of DNA replication and recovery from DNA damage.

To determine whether checkpoint suppression translated into a specific DNA break repair defect, we assessed both HR and non-homologous end joining (NHEJ) using the DR-GFP and EJ5-GFP reporter systems, respectively (Fig. 3g)^36,37^. FR054 progressively reduced HR-mediated repair in a concentration- and time-dependent manner. At 24 h, 500 μM FR054 decreased HR efficiency to approximately 65% of control levels. At 48 h, HR activity was reduced at both concentrations and dropped to approximately 55% at 500 μM (P = 0.020) (Fig. 3h). In contrast, NHEJ-mediated repair remained largely unaffected, with no significant reduction at either time points (Fig. 3i). Consistent with this defect in repair capacity, alkaline comet assays revealed increased DNA damage after all treatments, with damage remaining elevated upon co-treatment with GEM, particularly at 48 h (Fig. 3j, k). The comparable comet signal in the presence of checkpoint collapse and reduced HR suggests impaired processing or resolution rather than a further increase in initial lesion formation.

### FR054 reshapes the O-GlcNAc-proteome and suppresses GEM-associated GlcNAcylation programs

Beyond the canonical phosphorylation cascades activated by DNA damage, dynamic O-GlcNAcylation of chromatin-associated and DNA repair proteins provides an additional regulatory layer that modulates their recruitment, stability and activity during lesion recognition and repair. Because O-GlcNAcylation can cooperate with, fine-tune or counterbalance phosphorylation-dependent signaling^14,38,39^, we hypothesized that its reduction by FR054 could selectively weaken the repair response elicited by GEM. To investigate this mechanism, we selected MIAPaCa-2 cells because they displayed a robust and reproducible response to the combined treatment, characterized by checkpoint impairment, defective HR and persistent DNA damage. O-GlcNAcylated proteins were therefore immunoprecipitated from MIAPaCa-2 cells treated with FR054 or GEM, alone or in combination, for 48 h, and subsequently identified by LC-MS/MS. O-GlcNAcylated proteins were isolated through immuno-capture, trypsin-digested, and analyzed by nanoLC-MS/MS, followed by MaxQuant and Perseus processing (Fig. 4a and Supplementary Tables 6 and 7)^40,41^. Volcano-plot analysis revealed profound changes in the O-GlcNAc-enriched proteome in response to FR054. Compared with control cells, FR054 treatment caused a strong reduction in O-GlcNAcylated proteins, with 223 proteins significantly down-O-GlcNAcylated and only 41 upregulated (Fig. 4b, left panel). In contrast, GEM led to increased O-GlcNAcylation of 132 proteins, with far fewer downregulated targets (n=38), indicating that GEM activates rather than suppresses O-GlcNAcylation (Fig. 4b, middle panel). Remarkably, the combination reversed much of the GEM-induced O-GlcNAcylation, yielding a profile dominated by decreased O-GlcNAcylated proteins (148 down- versus 80 up-regulated; Fig. 4b, right panel). These data show that FR054 effectively counteracts the GEM-driven rise in O-GlcNAcylated proteins and broadly suppresses O-GlcNAcylation. Pathway enrichment using Ingenuity Pathway Analysis (IPA)^42^ identified major biological processes and cancer-relevant functions impacted by O-glycoproteome remodeling (Fig. 4c). Canonical pathway analysis predicted a broad inhibition of protein synthesis and RNA-processing programs, including ribosomal quality control, EIF2 signaling, translation initiation/elongation/termination, rRNA processing and nonsense-mediated decay. Pathways involved in DNA synthesis, cell-cycle checkpoints, G2/M progression and mitosis were also predicted to have reduced activity. Consistently, Diseases and Functions analysis predicted decreased cell survival, viability, proliferation, migration and invasion, together with increased apoptosis, cell death, cellular sensitivity and DNA damage. Overall, FR054 induces a strong anti-proliferative and cytotoxic remodeling of the O-GlcNAc- enriched proteome. In contrast, GEM predominantly increased the O-GlcNAcylation of proteins associated with translational control, RNA processing, amino-acid stress responses, energy metabolism, DNA synthesis and cell-cycle checkpoint pathways. Diseases and Functions analysis predicted activation of cell survival, viability, proliferation and motility, with a concomitant reduction in apoptosis and cell death. Thus, GEM appears to induce an adaptive O-GlcNAc-dependent program that may help pancreatic cancer cells tolerate replication stress. Strikingly, the combination largely reversed the GEM-induced profile, suppressing translational, metabolic, cell-cycle and DNA-replication pathways and reducing predicted survival, proliferation, migration and invasion. At the same time, apoptosis, cell death, cellular sensitivity and DNA damage functions were also predicted to have reduced activity. Overall, these analyses indicate that GEM induces a predominantly adaptive and pro-survival remodelling of the O-GlcNAc-enriched proteome, whereas FR054, alone and particularly in combination with GEM, suppresses translational and cell-cycle programmes and favors DNA damage and cell death.

**Figure 4.**
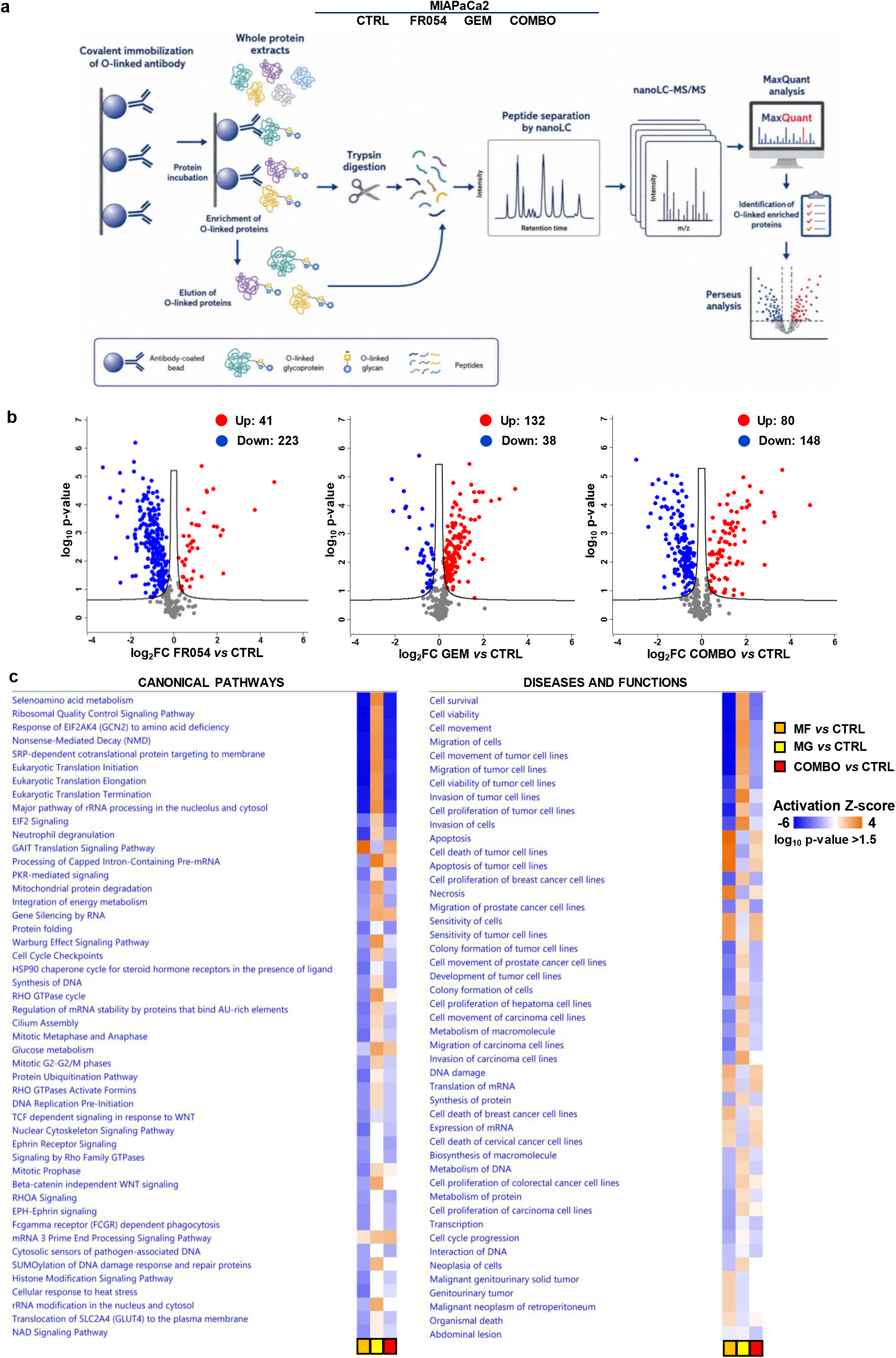
Mass spectrometry identifies GEM-induced O-GlcNAcylation of cancer-associated proteins and its suppression by FR054. **a,** Experimental workflow for the identification of O- GlcNAcylated proteins in MIAPaCa-2 cells treated for 48 h with vehicle (CTRL), 500 μM FR054, 1 μM gemcitabine (GEM), or their combination (COMBO). O-GlcNAcylated proteins were enriched by immunoprecipitation using the RL2 antibody and analysed by liquid chromatography-tandem mass spectrometry. Protein identification, quantification and differential enrichment analyses were performed using MaxQuant and Perseus. **b**, Volcano plots showing proteins differentially represented in the O-GlcNAc-enriched fraction following FR054 (left panel), GEM (middle panel) or COMBO (right panel) treatment relative to CTRL. FR054 treatment resulted in 41 increased and 223 decreased proteins, GEM treatment in 132 increased and 38 decreased proteins, and COMBO treatment in 80 increased and 148 decreased proteins. **c**, Ingenuity Pathway Analysis (IPA) of the differentially represented proteins, showing predicted changes in canonical pathways and disease and biological functions following each treatment relative to CTRL. Colour intensity indicates the predicted activation z-score, ranging from inhibition (blue) to activation (orange). Only categories with −log_10_ p- value > 1.5 are shown.

### GEM induces Thr81 O-GlcNAcylation of RUVBL2, stabilizing the RUVBL1-RUVBL2 complex without compromising ATP-Mg binding

Among the proteins showing treatment-dependent changes in O-GlcNAcylation, RUVBL2 emerged as a particularly relevant candidate. RUVBL2 forms an essential AAA+ ATPase complex with RUVBL1 and has recently been identified as a critical element in PDAC, where the RUVBL1/2 complex supports MYC-driven oncogenic transcription and KLF5-dependent classical and basal-like lineage programmes^43,44^. Moreover, the RUVBL1/2 complex has established roles in maintaining the stability and assembly of PIKK-containing complexes and in supporting chromatin remodelling, DNA replication, DNA repair and tolerance to replication-associated damage^19^. These functions made RUVBL2 a considerable candidate through which HBP-dependent O-GlcNAcylation could regulate the ability of pancreatic cancer cells to cope with GEM-induced DNA damage. LC-MS/MS identified multiple RUVBL2 peptides, including a glycopeptide carrying a HexNAc- modification on Thr81, a residue located within the Walker A nucleotide-binding motif of the AAA+ ATPase core^45^ (Fig. 5a-c and Extended Data Fig. 5a). Immunoprecipitation under denaturing conditions confirmed a strong increase in O-GlcNAcylated RUVBL2 specifically upon GEM treatment, indicating that this post-translational modification is dynamically regulated by genotoxic stress (Fig. 5d). To assess the evolutionary relevance of the modified residue, we performed a motif-focused alignment of the Walker A/P-loop region across representative eukaryotic RUVBL2 orthologues. The analysis showed that the threonine corresponding to human Thr81 is strictly conserved from vertebrates to yeast and plants, together with the catalytic lysine and the following threonine of the Walker A motif (Extended Data Fig. 5b). In contrast, the surrounding residues showed greater sequence variability. The selective conservation of Thr81 within an otherwise partially variable region supports the hypothesis that this residue has a functionally constrained role in eukaryotic RUVBL2 and may represent a conserved regulatory site within the ATP-binding pocket.

**Figure 5.**
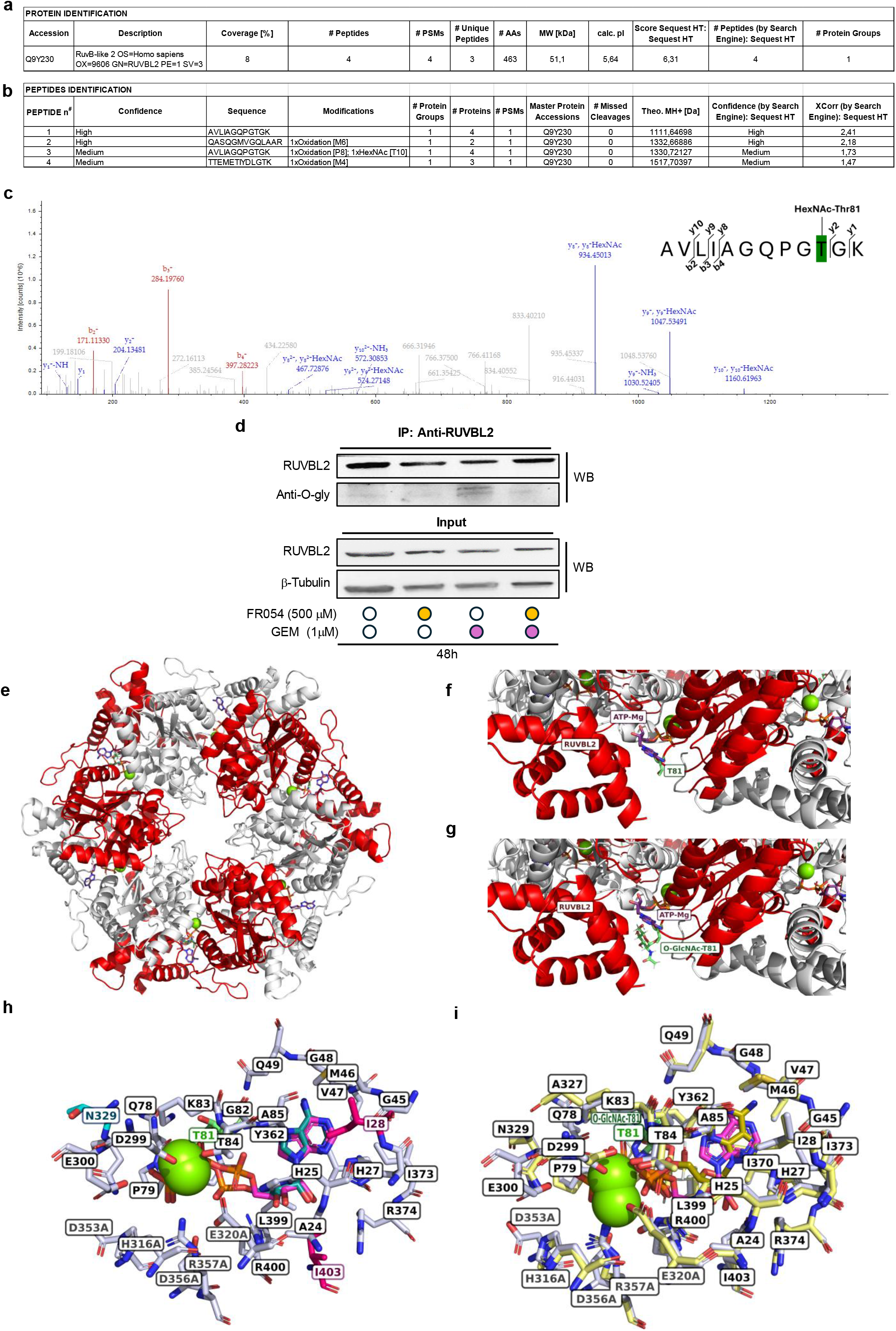
GEM promotes RUVBL2 O-GlcNAcylation, with Thr81 identified as an O-GlcNAc- modified residue. **a**, Proteome Discoverer output summarizing the identification of human RUVBL2 by liquid chromatography-tandem mass spectrometry (LC-MS/MS), including protein accession number, sequence coverage, number of identified and unique peptides, peptide- spectrum matches (PSMs), protein length, predicted molecular mass, calculated isoelectric point and Sequest HT score. **b**, Proteome Discoverer identification of RUVBL2-derived peptides following RUVBL2 immunoprecipitation, SDS-PAGE separation, in-gel tryptic digestion and LC-MS/MS analysis. The peptide AVLIAGQPGTGK was identified with a HexNAc modification on Thr81. Confidence scores, peptide sequences, post-translational modifications, PSMs, missed cleavages, theoretical peptide masses and Sequest HT scores are shown**. c**, Representative MS/MS fragmentation spectrum of the RUVBL2 glycopeptide AVLIAGQPGT*GK,* in which *T* corresponds to HexNAc-modified Thr81. The detected *b*- and *y*-fragment ions supporting peptide sequence assignment and HexNAc localization are indicated. **d**, RUVBL2 was immunoprecipitated from MIAPaCa-2 cells treated for 48 h with vehicle (CTRL), 500 μM FR054, 1 μM gemcitabine (GEM), or their combination (COMBO). Immunoprecipitated RUVBL2 was analysed by immunoblotting with antibodies against RUVBL2 and O-GlcNAc (RL2). RUVBL2 and β-tubulin were analysed in whole-cell lysates as input and loading controls, respectively. Immunoblots are representative of three independent biological experiments (*n*=3) with similar results. **e,** Top view of the computational model of the RUVBL1- RUVBL2 heterohexamer. RUVBL1 and RUVBL2 monomers are shown as white and red cartoons, respectively. ATP-Mg molecules occupying the nucleotide-binding sites are shown in magenta, and RUVBL2 Thr81 residues are highlighted in mint green. **f**, **g**, Enlarged side views of the ATP-Mg-binding region surrounding Thr81 in the non-glycosylated (**f**) and Thr81-O- GlcNAcylated (**g**) RUVBL1-RUVBL2 complexes. **h,** Comparison of the crystallographic and redocked ATP-Mg-binding poses in the non-glycosylated complex. The crystallographic and redocked ATP-Mg molecules are shown in magenta and sky blue, respectively. Residues located within 5 Å of both ATP-Mg poses are shown as white sticks, whereas residues found exclusively within 5 Å of the crystallographic or redocked pose are shown in dark pink and cyan, respectively. Distances involving residues unique to the crystallographic or redocked ATP-Mg-binding environment, together with the corresponding residue labels, are shown in magenta or blue, respectively; all other residue labels are shown in black. **i**, Comparison of the crystallographic and docked ATP-Mg-binding poses in the Thr81-O-GlcNAcylated complex. The crystallographic and docked ATP-Mg molecules are shown in magenta and gold, respectively. Residues located within 5 Å of the crystallographic ATP-Mg pose are shown as white sticks, whereas residues located within 5 Å of the docked pose in the Thr81-O- GlcNAcylated complex are shown as pale-yellow sticks. Grey and orange distance labels indicate interactions involving the crystallographic and docked ATP-Mg poses, respectively. The displayed residues and distances illustrate changes in the ATP-Mg-binding environment associated with Thr81 O-GlcNAcylation. E300 and N329 are highlighted in dark pink in **h**, **i** to show that their interactions with the ATP phosphate tail, which are present in the crystallographic structure and weakened following redocking in the non-glycosylated model, are restored in the docked glycosylated complex. In **h**, **i**, residue labels ending in A correspond to RUVBL1 chain A, whereas all other labelled residues belong to RUVBL2 chain D. Thr81 is highlighted in mint green.

Given the strategic location of Thr81, we investigated how its GlcNAcylation affects the structure and function of the RUVBL1-RUVBL2 hetero-hexamer^46^. Computational modeling revealed that Thr81 O-GlcNAcylation enhances the thermodynamic stability of the hexamer (Fig. 5e), lowering the Rosetta interface energy and increasing inter-subunit interactions (Supplementary Table 8a). The O-GlcNAcylated form displayed increased hydrogen bonding and a larger buried interface area, consistent with enhanced stabilization of the assembly (Fig. 5g and Supplementary Table 8a). ATP-Mg docking instead indicated that Thr81 O-GlcNAcylation does not compromise nucleotide binding. The non-glycosylated wild-type and Thr81-O-GlcNAcylated models showed similarly favorable predicted binding energies (−14.19 and −14.47 kcal/mol, respectively) and Ki values in the picomolar range (39.75 and 24.72 pM, respectively). Structural inspection indicated that O-GlcNAcylation induces subtle rearrangements within the ATP-binding pocket, involving residues such as E300 in the Walker B domain^47^ and N329, both important for ATP hydrolysis^48^ (Fig. 5h, i). These changes occur while maintaining a favorable predicted ATP-Mg-binding configuration. Collectively, these findings demonstrate that GEM induces O-GlcNAcylation of RUVBL2 at Thr81, a modification predicted to stabilize the RUVBL1-RUVBL2 heterohexamer while preserving favorable ATP-Mg engagement. This suggests that Thr81 O-GlcNAcylation may promote a structurally stable, nucleotide-competent RUVBL1-RUVBL2 assembly, with potential consequences for ATR/ATM biogenesis and stability and the DDR under chemotherapy-induced stress.

### HBP inhibition promotes nuclear accumulation of RUVBL2 in pancreatic cancer cells

In light of the established role of O-GlcNAcylation in regulating protein activity, stability and subcellular localization^7^, we next investigated whether the FR054-dependent reduction in RUVBL2 O-GlcNAcylation could affect its expression and intracellular distribution in MIAPaCa-2 and PANC-1 pancreatic cancer cells. Immunoblotting showed that total RUVBL2 protein abundance remained largely unchanged across treatment conditions in both cell lines (Fig. 6a-d), indicating that neither FR054 nor GEM substantially affected RUVBL2 expression. Immunofluorescence analysis, however, revealed marked and opposing effects on RUVBL2 subcellular distribution. FR054 significantly increased the nuclear-to-cytoplasmic ratio of RUVBL2, consistent with enhanced nuclear accumulation (Fig. 6e-h). In contrast, GEM reduced this ratio relative to the corresponding control, indicating preferential redistribution or retention of RUVBL2 in the cytoplasmic compartment. This effect became more pronounced at 48 h (Fig. 6e-h). Notably, combined treatment counteracted the GEM-induced cytoplasmic shift and restored RUVBL2 nuclear accumulation, with a stronger effect in MIAPaCa-2 cells and a partial recovery in PANC-1 cells.

**Figure 6.**
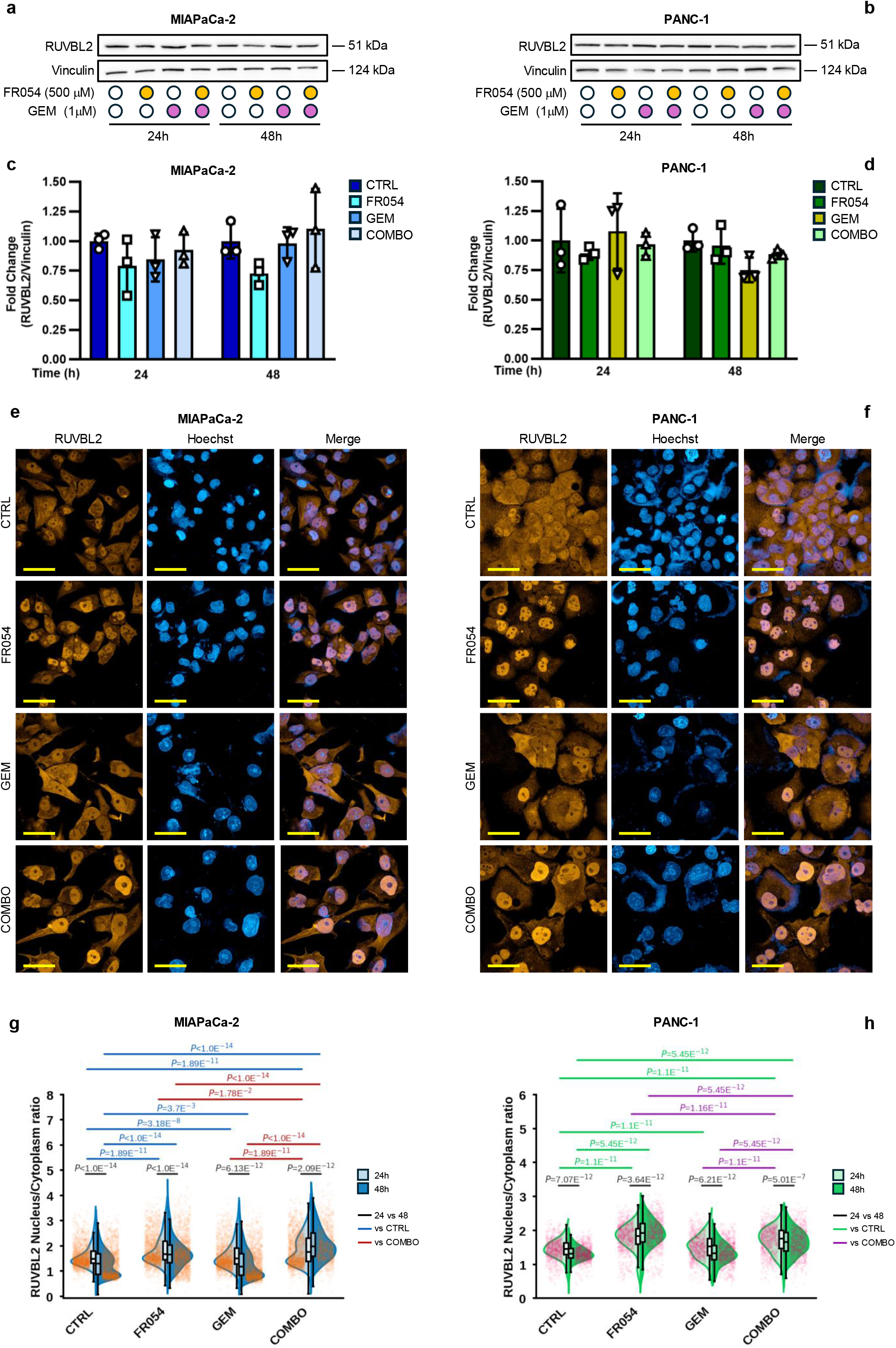
PGM3 inhibition promotes the nuclear redistribution of RUVBL2 without affecting its overall abundance. **a, b**, Immunoblot analysis of RUVBL2 in MIAPaCa-2 (**a**) and PANC-1 (**b**) cells treated with vehicle (CTRL), 500 μM FR054, 1 μM gemcitabine (GEM), or their combination (COMBO) for 24 h or 48 h. Vinculin was used as a loading control. Representative data from three independent experiments are shown (*n*=3). **c, d**, Densitometric quantification of RUVBL2 protein abundance in MIAPaCa-2 (**c**) and PANC-1 (**d**) cells. RUVBL2 signals were normalized to vinculin and expressed as fold change relative to the corresponding time-matched CTRL (*n*=3 independent biological experiments). **e, f**, Representative immunofluorescence images showing the subcellular distribution of RUVBL2 in MIAPaCa-2 (**e**) and PANC-1 (**f**) cells following treatment with CTRL, FR054, GEM or COMBO. RUVBL2 is shown in red and nuclei were counterstained with Hoechst (blue). Scale bar, 50 μm (63× magnification). **g, h**, Single-cell quantification by Harmony software of the nuclear-to- cytoplasmic RUVBL2 fluorescence intensity ratio in MIAPaCa-2 (**g**) and PANC-1 (**h**) cells after 24 h and 48 h of treatment as described in **a**, **b**. Split violin plots show the distributions at 24 h and 48 h, and each point represents an individual cell. Embedded box plots indicate the median, interquartile range (IQR; box limits), and 1.5×IQR whiskers. Black annotations indicate comparisons between 24 h and 48 h within each treatment; blue and green annotations indicate comparisons with the corresponding CTRL condition in MIAPaCa-2 (**g**) and PANC-1 (**h**) cells, respectively; red (**g**) and purple (**h**) annotations indicate comparisons with the corresponding COMBO condition. *n*=2742–22519 (**g**) and *n*=1549–5272 (**h**) cells were analysed per condition from at least three biological replicates (*n*≥3). In **c** and **d**, data are presented as mean ± s.d. Statistical significance was determined using two-way ANOVA with Tukey’s multiple comparisons test (**c**, **d**, **g** and **h**). Exact *P* values are reported in the plots; ns, not significant.

These findings indicate that the treatments primarily affect RUVBL2 intracellular trafficking rather than its overall expression, with HBP inhibition opposing the cytoplasmic redistribution induced by GEM. Together with the glycoproteomic and structural data, these results suggest that GEM-induced O-GlcNAcylation and HBP inhibition exert opposing effects on RUVBL2 nucleocytoplasmic distribution, potentially redirecting its activity toward distinct subcellular functions involved in ATR/ATM protein homeostasis and the DDR.

### Thr81 O-GlcNAcylation regulates RUVBL2 protein localization, ATR/ATM expression and response to GEM

Because Thr81 lies within the Walker A nucleotide-binding motif of RUVBL2 (Extended Data Fig. 5a), a region central to ATP engagement and nucleotide-dependent conformational control, modification of this residue would be expected to have direct consequences for RUVBL2 function. To investigate whether Thr81 O-GlcNAcylation affects RUVBL2 function during genotoxic stress, we generated a Threonine 81 to Alanine mutant RUVBL2 (T81A), which removes the candidate O-GlcNAc acceptor residue (Extended Data Fig. 7a). Computational structural modelling of the ATP-binding pocket indicated that the T81A substitution alters the local organization of residues interacting with ATP-Mg and substantially weakens predicted nucleotide binding compared with both the non-glycosylated wild-type and Thr81-O-GlcNAcylated structures (Extended Data Fig. 7a and Supplementary Table 8b).

We then examined the functional consequences of this mutation in HEK293T cells transiently transfected with a construct co-expressing either the GFP and HA-RUVBL2 WT or the GFP and HA-RUVBL2-T81A (RUVBL2 MUT) (Extended Data Fig. 7b). Immunofluorescence analysis of HA-tagged RUVBL2 showed that the T81A substitution was associated with altered subcellular distribution relative to the wild-type protein (Fig. 7a, b). Quantification of the nuclear-to-cytoplasmic HA signal indicated that, in both untreated and GEM-treated cells, the T81A mutation promoted nuclear enrichment of RUVBL2 (Fig. 7b), consistent with the role for Thr81 and its O-GlcNAcylation in regulating RUVBL2 intracellular dynamics, as also suggested by the effects observed following FR054 treatment (Fig. 6e-h). Because RUVBL-containing complexes have been implicated in the regulation of phosphatidylinositol 3-kinase-related kinase (PIKK) family members^19,49^, we next assessed ATR and ATM protein levels in cells expressing WT or mutant RUVBL2 proteins. Western blot analysis showed that RUVBL2 MUT markedly reduced total ATR abundance under basal conditions and prevented its maintenance or accumulation following GEM treatment (Fig. 7c, d). Total ATM levels showed a similar, although less pronounced, pattern. Most ATM comparisons did not reach statistical significance; nevertheless, RUVBL2 MUT consistently reduced ATM abundance in both untreated and GEM-treated cells, with the difference reaching significance compared with NO DNA sample after GEM exposure (Fig. 7c, e). Thus, mutation of Thr81 appears to exert a stronger effect on ATR expression, while also showing a reproducible trend toward reduced expression of ATM (Fig. 7c-e). In order to define the effect of the mutant upon GEM treatment, HEK293T- transfected cells as before were monitored by live-cell imaging for 72 h post-transfection of which 48 h of GEM treatment (Extended Data Fig. 7c). Incucyte live-cell imaging revealed distinct confluence kinetics between WT and T81A RUVBL2-expressing cells. Although GEM-treated cells appeared to reach apparent confluence earlier at intermediate time points, likely due to treatment-induced increase in nuclei size (Fig. 7a), the overall confluence profile was reduced in T81A-expressing cells compared with WT controls (Fig 7f, g). These data are consistent with increased sensitivity of the T81A mutant to GEM effects. Overall, these findings suggest that Thr81 O-GlcNAcylation contributes to RUVBL2 cytoplasmic localization and maintenance of ATR and ATM protein levels and that loss of this O-GlcNAcylation site increases cellular sensitivity to GEM.

**Figure 7.**
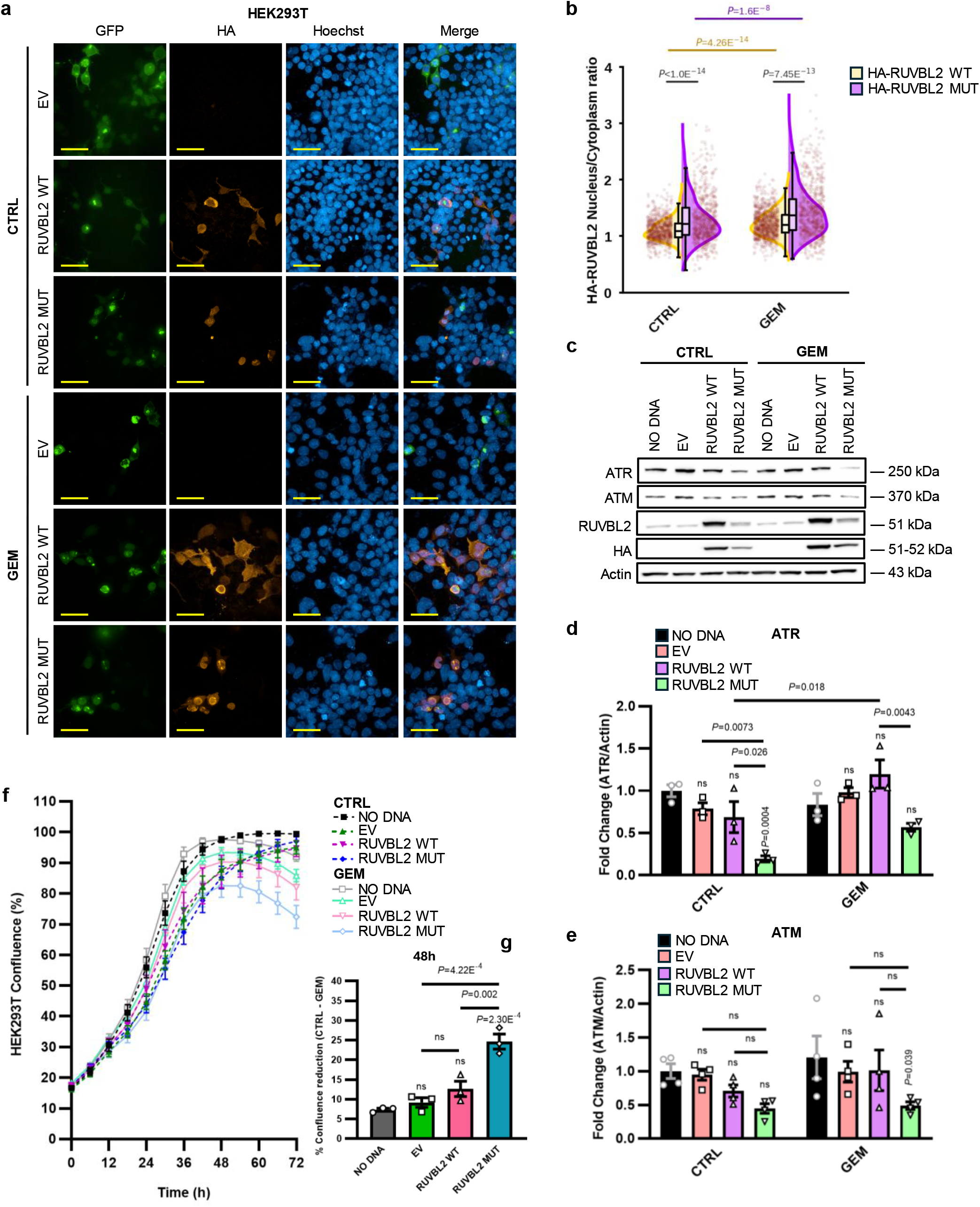
RUVBL2 Thr81 mutation alters RUVBL2 subcellular localization, ATR and ATM abundance and the response to GEM. **a**, Representative immunofluorescence microscopy images of HEK293T cells transfected with empty vector (EV) or constructs encoding HA-tagged wild- type RUVBL2 (RUVBL2 WT) or the RUVBL2 T81A mutant (RUVBL2 MUT) and treated with vehicle (CTRL) or 1 μM gemcitabine (GEM). GFP fluorescence identifies transfected cells, HA immunofluorescence detects ectopically expressed RUVBL2 (red), and nuclei were counterstained with Hoechst (blue). Images are representative of three independent experiments with similar results (*n*=3); scale bar, 50 μm (63× magnification). **b,** Single-cell quantification by Harmony software of the nuclear-to-cytoplasmic HA-RUVBL2 fluorescence- intensity ratio in cells expressing RUVBL2 WT or RUVBL2 MUT under CTRL or GEM treatment. Split violin plots show the distribution of individual cells expressing RUVBL2 WT or RUVBL2 MUT (*n*=692–1228 cells per condition from *n*=3 biological replicates). Embedded box plots indicate the median, interquartile range (IQR; box limits), and 1.5×IQR whiskers. **c,** Immunoblot analysis of ATR, ATM, total RUVBL2 and HA in non-transfected cells (NO DNA) and cells transfected with EV, RUVBL2 WT or RUVBL2 MUT, in the absence or presence of GEM. Representative immunoblots from *n*≥3 independent biological experiments are shown, and actin was used as a loading control. **d**, **e**, Densitometric quantification of ATR (**d**) and ATM (**e**) abundance. Protein signals were normalized to actin and expressed as fold change relative to the NO DNA-CTRL (ATR, *n*=3; ATM, *n*=4 independent biological experiments). **f,** Real-time monitoring of HEK293T cell confluence over 72 h using the Incucyte Live-Cell Analysis System. Confluence analysis was performed in non-transfected HEK293T cells and in cells expressing EV, RUVBL2 WT, or RUVBL2 MUT, cultured in the absence or presence of GEM. Data represent the analysis of *n*=15 fields per condition at 10× magnification from *n*=3 independent biological experiments. **g,** Difference in cell confluence between CTRL- and GEM- treated cultures at 48 h, showing an increased response to GEM in cells expressing RUVBL2 MUT (*n*=3 independent biological experiments). Data in **d**, **e** and **g** are presented as mean ± s.e.m. Statistical significance was determined using two-way ANOVA with Tukey’s multiple comparisons test (**b**, **d**, and **e**) or one-way ANOVA with Tukey’s multiple comparisons test (**g**). Exact *P* values are indicated in the plots; ns, not significant.

### FR054 sensitizes pancreatic cancer cells to treatment with olaparib and ionizing radiation by amplifying DNA damage

Because FR054 disrupts ATR/ATM signaling and impairs HR, thereby promoting an HR-deficient, BRCAness-like state, we next tested whether HBP inhibition could enhance sensitivity to the PARP inhibitor olaparib, which preferentially targets HR-deficient cells^50–52^, and to γ-irradiation, which induces DNA double-strand breaks whose error-free repair relies on functional HR^53,54^. Bliss independence analysis revealed a synergistic interaction between FR054 and olaparib in both pancreatic cancer cell lines, with maximum Bliss scores of 27 in MIAPaCa- 2 and 28 in PANC-1 cells (Extended Data Fig. 8a, b). For subsequent mechanistic experiments, we selected 350 μM FR054 and 20 μM olaparib. This combination maintained the FR054 concentration used throughout the study, showed synergistic activity in both cell lines and allowed direct comparison of their molecular responses. Consistent with the Bliss analysis, the combination increased cell death in both models, with the strongest effect observed at 72 h and a more rapid response in PANC-1 cells (Extended Data Fig. 8c, d).

To determine whether the increased cytotoxicity of the FR054-olaparib combination was associated with defective HR repair, we examined γH2AX accumulation and BRCA1 recruitment to nuclear foci, using total ATM protein abundance as an additional readout of DNA damage signaling competence. At 24 h, FR054 increased γH2AX foci in MIAPaCa-2 cells, both alone and in combination with olaparib, while markedly reducing BRCA1 focal accumulation (Extended Data Fig. 8e, g). Olaparib alone induced a more limited increase in γH2AX and largely preserved BRCA1 recruitment (Extended data Fig. 8e, g). In PANC-1 cells, FR054 and the combination also increased γH2AX, whereas BRCA1 foci were reduced, with the strongest decrease observed following combined treatment (Extended Data Fig. 8f, h). Thus, in both models, HBP inhibition promoted DNA damage accumulation without a proportional recruitment of BRCA1 to damaged chromatin. Strikingly, at 48 h, the dissociation between γH2AX accumulation and BRCA1 recruitment became more evident. In MIAPaCa-2 cells, the combination produced a marked increase in γH2AX foci together with a pronounced reduction in BRCA1 focal localization (Fig. 8a, c). FR054 alone showed a similar, although less extensive, pattern (Fig. 8a, c). In PANC-1 cells, γH2AX remained elevated following FR054 and combined treatment, while BRCA1 recruitment remained reduced, although the effect was more variable than in MIAPaCa-2 cells (Fig. 8b, d). These findings indicate that HBP inhibition impairs the spatial recruitment of BRCA1 despite persistent DNA damage, consistent with defective engagement of the HR machinery. Consistent with this phenotype, FR054 also reduced total ATM protein abundance. In both MIAPaCa-2 and PANC-1 cells, FR054 induced a similar time-dependent reduction in total ATM abundance. At 24 h, ATM levels showed a downward trend that did not reach statistical significance (Fig 8e, f and Extended Data 8i, j), whereas at 48 h the decrease became more pronounced and statistically significant (Fig 8e, f and Extended Data 8i, j). This effect was also maintained in the combination with olaparib, while olaparib alone did not reproduce the same reduction (Fig 8e, f and Extended Data 8i, j). These findings indicate that HBP inhibition progressively compromises the maintenance of ATM protein levels in both pancreatic cancer cell lines. Collectively, these data show that FR054 increases DNA damage burden while reducing BRCA1 recruitment and ATM abundance, thereby creating a cellular context compatible with impaired HR repair and enhanced sensitivity to olaparib.

**Figure 8.**
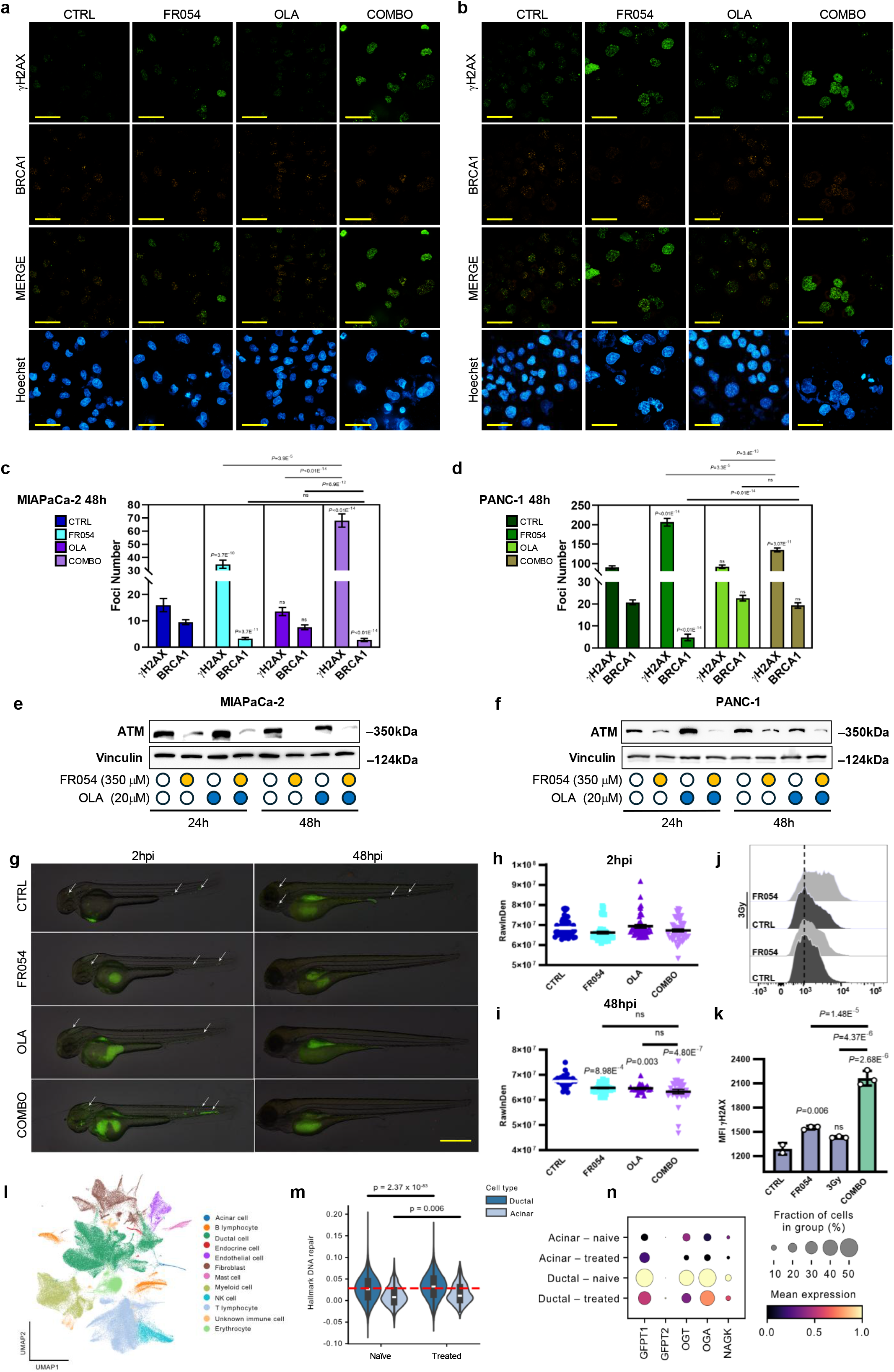
PGM3 inhibition impairs HR DNA repair response and potentiates the effects of olaparib and ionizing radiation in PDAC. **a**, **b**, Representative immunofluorescence images of γH2AX and BRCA1 in MIAPaCa-2 (**a**) and PANC-1 (**b**) cells treated for 48 h with vehicle (CTRL), 350 μM FR054, 20 μM olaparib (OLA), or their combination (COMBO). γH2AX is shown in green, BRCA1 in red and nuclei were counterstained with Hoechst (blue). Images are representative of three independent experiments with similar results (*n*=3); scale bar, 50 μm (63× magnification). **c**, **d**, Quantification of γH2AX and BRCA1 nuclear foci in MIAPaCa-2 (**c**) and PANC-1 (**d**) cells under the conditions shown in **a**, **b**. Bars represent the mean values obtained from *n*=98–177 (**c**) or *n*=150–277 (**d**) nuclei per condition, pooled from three independent biological experiments. **e**, **f**, Immunoblot analysis of ATM in MIAPaCa-2 (**e**) and PANC-1 (**f**) cells treated with CTRL, FR054, OLA or COMBO for 24 h or 48 h. Vinculin was used as a loading control. Representative data from three independent experiments with similar results are shown (*n*=3). **g**, Representative fluorescence images from three independent biological experiments showing zebrafish embryos bearing GFP-labelled PDAC xenografts under the indicated treatment conditions, acquired at 2 h and 48 h post-injection (hpi). Arrows indicate disseminated tumor cells. Scale bar, 200 μm. **h**, **i**, Quantification of tumor- associated GFP fluorescence, expressed as raw integrated density (RawIntDen), at 2 hpi (**h**) and 48 hpi (**i**). Each symbol represents an individual embryo (2 hpi: *n*=49 for each condition; 48 hpi: CTRL, *n*=37; FR054, *n*=39; OLA, *n*=22; COMBO, *n*=32 embryos) from *n*=3 biological experiments. No statistically significant differences in tumor-associated GFP fluorescence were observed at 2 hpi among the different treatment conditions, whereas changes detected at 48 hpi reflect treatment-dependent effects on tumor growth. **j**, Representative flow-cytometry histograms from at least two independent biological replicates showing γH2AX fluorescence in cells treated with CTRL or FR054 and either left unirradiated or exposed to 3 Gy. **k**, Quantification of γH2AX mean fluorescence intensity (MFI) in CTRL, FR054-treated, 3-Gy- irradiated and combined FR054-plus-irradiation (COMBO) conditions. Data were obtained from at least two independent biological replicates (*n*≥2). **l**, Uniform manifold approximation and projection (UMAP) representation of single-cell transcriptomes from treatment-naive and treated PDAC samples, colored according to the annotated cell populations. **m**, Distribution of the Hallmark DNA repair signature score in acinar and ductal cells from treatment-naive and treated samples. Boxes indicate the median and interquartile range. **n**, Dot plot showing the expression of the HBP-associated genes *GFPT1*, *GFPT2*, *OGT*, *OGA* and *NAGK* in acinar and ductal cells from treatment-naive and treated samples. Dot size represents the fraction of cells expressing each gene, and color indicates scaled mean expression. Data are presented as mean ± s.e.m. (**c**, **d**, **h** and **i**) or as mean ± s.d. (**k**). Statistical significance was determined using Kruskal-Wallis test followed by Dunn’s post-hoc test (**c** and **d**) and one-way ANOVA followed by Bonferroni’s post hoc test (**h** and **i**) or Tukey’s multiple comparisons test (**k**). Cell-level statistical comparisons of gene set activity between untreated and treated ductal and acinar cells were performed using a two-sided Wilcoxon rank-sum test (**m**). Exact *P* values are indicated in the plots; ns, not significant.

To validate these findings in vivo, we used a zebrafish xenograft model, a rapid and reliable platform for assessing short-term tumor-cell persistence and drug response. MIAPaCa-2 cells were FITC-labelled and treated in vitro for 24 h with FR054, olaparib or their combination before injection into zebrafish larvae. Tumor-associated fluorescence was comparable among groups at 2 h post-injection (Fig 8g, h) indicating similar initial engraftment. At 48 h post- injection, all treatments significantly reduced tumor-cell burden compared with the control (Fig 8g, i). The combination produced the lowest tumor-associated fluorescence, consistent with reduced persistence of pretreated tumor cells after engraftment.

Because HR-deficient cells are less able to accurately repair DNA double-strand breaks, induction of a BRCAness-like state is expected to increase sensitivity to ionizing radiation. We therefore first characterized the γH2AX response of MIAPaCa-2 cells to γ-irradiation. Exposure to 1, 3 or 5 Gy induced a rapid and dose-dependent increase in γH2AX fluorescence, detectable at the earliest time points and persisting for up to 48 h, particularly at the higher doses (Supplementary Fig. 2a). The signal then declined between 48 h and 72 h, suggesting partial recovery and progressive resolution of irradiation-induced DNA damage in the surviving cell population. FR054 alone also increased γH2AX in a concentration-dependent manner, further supporting the ability of HBP inhibition to promote DNA damage accumulation and providing the rationale for testing its interaction with γ-irradiation (Supplementary Fig. 2b). We next asked whether FR054 could enhance the response to irradiation. Although both treatments individually increased γH2AX at 48 h, the combination produced the largest shift in fluorescence intensity and the highest γH2AX signal (Fig. 8j, k). This finding indicates that FR054 increases the persistence of irradiation-induced DNA damage, consistent with impaired DSB resolution in cells driven toward an HR-deficient, BRCAness-like state. To evaluate the clinical relevance of our findings, we next asked whether HBP-related metabolic features and DNA repair programs remain associated in human PDAC, where metabolic rewiring and DDR proficiency contribute to therapeutic resistance^55,56^. We reanalyzed publicly available single-cell RNA-sequencing data from treatment-naïve and chemotherapy-treated pancreatic cancer specimens^57^. Cell populations were annotated according to established lineage markers (Fig. 8l and Supplementary Table 9). DNA repair activity was quantified using the 150-gene MSigDB Hallmark DNA Repair signature listed in Supplementary Table 10. Acinar-like cells, representing a more differentiated epithelial state, displayed lower DNA repair activity than ductal tumor cells, with a further reduction following chemotherapy (Fig. 8m). This decrease was accompanied by lower expression of HBP-related genes, including GFPT1, GFPT2, OGT, OGA and NAGK (Fig. 8n). By contrast, malignant ductal cells retained comparatively higher DNA repair scores and HBP-associated gene expression in both treatment-naïve and treated samples, supporting a potential role for these programs in tumor-cell fitness and therapeutic resistance.

## Discussion

Pancreatic ductal adenocarcinoma remains one of the most therapeutically refractory malignancies, in part because of its marked ability to tolerate the replication stress and DNA damage induced by standard treatments such as GEM. Increasing evidence indicates that metabolic rewiring contributes to this adaptability, and our findings identify inhibition of HBP flux by the PGM3 inhibitor FR054 as a strategy to weaken DDR capacity and sensitize pancreatic cancer cells to GEM, PARP inhibition and ionizing radiation.

Consistent with previous evidence that O-GlcNAcylation regulates DDR signaling and influences DBS repair-pathway choice^12^, we found that FR054 suppressed global O-GlcNAcylation and compromised both ATR- and ATM-dependent signaling. In line with reports showing that therapy-induced O-GlcNAcylation enhances DSB repair and protects cancer cells from therapy-induced senescence^58^, HBP blockade prevented efficient checkpoint engagement, promoted replication stress and preferentially impaired HR while largely preserving NHEJ. This shift toward an HR-deficient state provides a mechanistic explanation for the persistence of DNA damage and the marked cytotoxic interaction between FR054 and GEM.

Our findings further identify RUVBL2 as a stress-responsive O-GlcNAcylated protein and suggest that Thr81 contributes to the cellular response to GEM. Thr81 lies within the Walker A nucleotide-binding motif and is therefore well positioned to influence RUVBL2 function. GEM increased RUVBL2 O-GlcNAcylation, with Thr81 identified as a treatment-regulated modification site by glycoproteomic analysis. Structural modelling predicted that Thr81 O-GlcNAcylation stabilizes the RUVBL1–RUVBL2 hexameric complex without compromising ATP-Mg binding, whereas the T81A substitution substantially weakened predicted nucleotide engagement. These findings are consistent with a model in which Thr81 O-GlcNAcylation favors a stable, nucleotide-competent RUVBL1-RUVBL2 assembly. This interpretation is compatible with the established function of RUVBL1/RUVBL2 as ATP-dependent assembly factors and scaffolds within large molecular complexes, including INO80, SRCAP and the R2TP/PAQosome, and with their requirement for the maturation and stability of PIKK family members^59–64^.

In parallel, FR054 promoted nuclear accumulation of RUVBL2 without substantially altering total protein abundance, indicating that perturbation of HBP/O-GlcNAcylation metabolism affects not only its biochemical state but also its subcellular distribution. This is consistent with previous evidence that RUVBL proteins display dynamic nucleocytoplasmic localization and can redistribute to specialized compartments, including the nucleolus and the midbody, depending on the cellular context^65–67^.

To our knowledge, these findings provide the first direct link between RUVBL2 O-GlcNAcylation and HBP flux and offer a possible mechanistic basis for previous observations connecting RUVBL-containing complexes to nutrient availability. Glucose and glutamine regulate the TTT-RUVBL complex involved in mTORC1 signaling, while RUVBL2 has also been linked to insulin-stimulated glucose uptake and broader metabolic control^68–71^. Notably, the TTT-RUVBL complex contributes to the assembly and stability of PIKK-containing complexes^18,19^, further supporting a functional connection between nutrient metabolism and the DNA repair machinery. Accordingly, the increased GEM sensitivity and reduced ATR/ATM abundance associated with the T81A mutant are consistent with a role for Thr81, and potentially its O-GlcNAcylation, in preserving RUVBL2-dependent complex function and PIKK homeostasis during genotoxic stress.

This interpretation also fits with the higher expression^72^ and wider role of RUVBL proteins in cancer. RUVBL1/2 support tumor-cell proliferation, DNA replication, stress tolerance and oncogenic signaling, while pharmacological inhibition of their ATPase activity impairs cancer-cell fitness in several tumors, including lung cancer and mTORC1-hyperactivated cancers^19,49,73^. The allosteric RUVBL1/2 inhibitor CB-6644 shows anticancer activity and reduces tumor growth in preclinical models^73^. RUVBL2 has also been implicated in tumor maintenance in acute myeloid leukaemia and hepatocellular carcinoma, where its depletion compromises proliferation, survival and DDR^74–76^.

In this context, induction of a BRCAness-like state by HBP inhibition is particularly relevant in PDAC, where HR defects create an actionable vulnerability to PARP inhibitors and other DNA-damaging therapies. This is illustrated by the clinical activity of olaparib in BRCA-mutated pancreatic cancer and by ongoing efforts to extend this strategy to HR-deficient tumors lacking canonical BRCA alterations^52,77^ Accordingly, the enhanced activity of the FR054-olaparib combination is mechanistically consistent with the ability of FR054 to induce an HR-deficient state. Increased γH2AX, an elevated γH2AX-to-BRCA1 foci ratio and persistent DNA damage under combined treatment all indicate defective lesion resolution.

The same model may explain the enhanced accumulation and persistence of radiation-induced DNA damage following FR054 treatment, because impaired ATM signaling is expected to compromise recovery from irradiation-induced lesions^78^. These findings are particularly relevant in PDAC, a tumor type characterized by sustained dependence on DNA repair programs. Consistent with this interpretation, our single-cell transcriptomic reanalysis showed that malignant ductal cells maintain higher DNA repair and HBP-associated activity than differentiated acinar cells^57^, supporting the idea that HBP inhibition exposes a repair dependency in pancreatic cancer. This effect was also observed in the zebrafish xenograft model, in which the combination produced the lowest tumor-associated fluorescence, consistent with enhanced suppression of tumor-cell persistence compared with either monotherapy^79^.

Overall, our findings support a model in which HBP inhibition induces a therapy-dependent BRCAness-like phenotype, at least partly through altered RUVBL2 function, reduced PIKK abundance, and impaired DDR-complex activity. This state sensitizes pancreatic cancer cells to PARP inhibition and enhances the persistence of radiation-induced DNA damage, highlighting HBP/O-GlcNAcylation signaling as a potentially actionable metabolic vulnerability in PDAC. Whether the HBP-RUVBL2-DDR axis identified here represents a broader vulnerability of HBP-dependent tumors remains to be established^80^. Nevertheless, the growing evidence that HBP activation supports tumor-cell survival, metabolic adaptation and resistance to anticancer therapies across breast, pancreatic, KRAS/LKB1-mutant lung cancer and glioblastoma^11,15,16,81,82^ suggests that this pathway may represent a broader determinant of therapeutic resistance. These observations support further investigation of HBP dependency across additional tumor types and treatment contexts.

## Methods

### Cell lines and tissue culture

Human pancreatic ductal adenocarcinoma cell lines MIAPaCa-2, PANC-1 and HEK293T were routinely cultured in high-glucose Dulbecco’s medium Eagle’s medium (DMEM) supplemented with 2 mM L-glutamine, 100 U/ml penicillin, 100 μg/ml streptomycin, and 10% fetal bovine serum (FBS). The cells were grown and maintained according to standard cell culture protocols and kept at 37°C with 5% CO_2_. The medium was replaced every 2-3 days and cells were split or seeded for experiments when they reached the sub-confluence. Cells were originally obtained from the American Type Culture Collection (ATCC).

### Trypan blue vital assay

Where not differently specified, for experiments, cells were seeded in the complete growth medium, and after 24 h, cells were washed twice with phosphate buffer saline 1X (PBS 1X) and incubated in a complete medium containing the indicated treatments. The main experimental conditions included treatment with 1 μM GEM and/or 500 μM FR054, or with 20 μM olaparib and/or 350 μM FR054, depending on the experimental setting. Treatments were maintained for the indicated time intervals. GEM and olaparib were purchased from Sigma-Aldrich, while FR054 was synthesized in collaboration with the laboratory of Prof. Barbara La Ferla at the University of Milano-Bicocca^15,83^.

Trypan blue vital assay was performed by seeded 1.25 × 10^5^ (MIAPaCa-2 and PANC-1) viable cells per well in 6-well plate. Cell death and cell viability were assessed at different time points 24 h, 48 h and 72 h, counting harvested cells with Burker chamber after staining with 0.4% trypan blue.

### Western blot analysis

MIAPaCa-2 and PANC-1 were seeded at 7.50 × 10^5^ onto 100-mm dishes in the complete growth medium. HEK293T cells were seeded in 6-well plate at a density of 8 × 10^4^ cells per well. After 24 h and 48 h of treatments, PDAC cells were harvested, whereas HEK293T cells were harvested 48 h after transfection, with GEM added during the final 24 h. Cells were disrupted in a buffer containing 50 mM Tris-HCl, pH 7.6, 150 mM NaCl, 1% (v/v) Triton X-100, 0.2% (v/v) sodium dodecyl sulfate (SDS), 0.5% (v/v) sodium deoxycholate, 1 mM MgCl2, 1 mM EDTA, protease inhibitor cocktail, and phosphatase inhibitors; 30 μg of total proteins was resolved by SDS-PAGE and transferred to the nitrocellulose membrane, which was incubated overnight with specific primary antibodies: γH2AX (#9718; 1:1000), cleaved caspase-3 (#9662S; 1:500), phospho-RPA32 Ser8 (#54762; 1:1000), RPA32 (#52448; 1:1000), phospho-CHK1 Ser345 (#2348; 1:1000), phospho-CHK2 Thr68 (#2661; 1:1000) from Cell Signaling Technology Inc.; Cyclin D1 (sc-20044; 1:1000),Cyclin E (sc-481; 1:200), Cyclin A (sc-271682; 1:500), Cyclin B1 (sc-245; 1:1000), ATR (sc-515173; 1:100), phospho-ATM Ser1981 (sc-47739; 1:500), ATM (sc- 377293; 1:100) from Santa Cruz Biotechnology Inc.; phospho-ATR Thr-1989 (GTX128145; 1:500) from GENETEX, RUVBL2 (HPA067966; 1:10.000), β-actin (A54411;10.000) from Sigma-Aldrich, Merck Life Science; HA (AM20783PU-N; 1:800) from Life Technologies-ThermoFisher Scientific; vinculin (#MAB80145; 1:10000) and β-tubulin (MAB-80143; 1:5000) from Immunological Sciences. Levels of protein expression on Western blots were quantified by densitometry analysis using ImageLab software. Confocal fluorescence microscopy For immunofluorescence detection, MIAPaCa-2, PANC-1 and HEK239T cells were seeded in complete growth medium in 96-well plates at a density of 5 × 10^3^ cells per well. PDAC cells were treated for 24 h or 48 h, whereas HEK293T cells were transfected for 48 h and exposed to GEM during the final 24 h. Cells were fixed with 4% paraformaldehyde for 10 min at room temperature (RT), followed by permeabilization with 0.3% Triton X-100 for 5 min. Subsequently, nonspecific binding sites were blocked using 1% BSA in PBS 1X for 1 h at RT. Cells were then incubated at 37 °C with the following primary antibodies: γH2AX (#9718, Cell Signaling Technology Inc.; 1:400); Cyclin A (sc-271682; 1:200), γH2AX Ser139 (sc-517348 1:400) and BRCA1 (sc-6954; 1:100) from Santa Cruz Biotechnology Inc.; HA (AM20783PU-N, Life Technologies-ThermoFisher Scientific; 1:800). After 1 h, cells were washed twice and incubated for 1 h at 37 °C with appropriate conjugated secondary antibodies: Alexa Fluor 488- conjugated secondary antibodies goat anti-mouse IgG (IS-20018, 1:500), Alexa Fluor 546- conjugated secondary antibodies goat anti-mouse IgG (IS13202, 1:500) from Immunological Sciences or Alexa Fluor 488-conjugated secondary antibodies donckey anti-rabbit IgG (A- 21206, Life Technologies-ThermoFisher Scientific; 1:500). Finally, nuclei were counterstained with Hoechst (2 μg/mL) for 10 min at RT. Fluorescence images were acquired using an Operetta CLS High-Content Analysis System (Revvity) with water-immersion 40× or 63× magnification and analyzed by Harmony software.

### EdU-Click assay

For the detection of proliferating cells, EdU (Click-iT, IS-873A, Immunological Sciences) incorporation was performed 1 h before fixation. MIAPaCa-2 and PANC-1 cells cultured in 96- well plates at the density of 5 × 10^3^ cells per well and treated for 6 h, 24 h or 48 h were incubated with 10 μM EdU for 1 h at 37 °C in complete medium. After incubation, wells were washed once with PBS 1X and fixed with 4% paraformaldehyde for 15 min at RT. Cells were then permeabilized with 0.5% Triton X-100 in PBS for 20 min, followed by two washes with 3% BSA in PBS 1X. The Click-iT reaction was performed according to the manufacturer’s protocol. Wells were incubated for 30 min at RT, followed by nuclear staining with Hoechst (2 μg/mL) for 10 min at RT. Fluorescence images were acquired using Operetta CLS High-Content Analysis System (Revvity) with water-immersion 40× magnification and analyzed by Harmony software.

### Double strand break (DSB) repair reporter assay

HR and NHEJ efficiencies were measured using previously established DR-GFP and EJ5-GFP U2OS cell lines, respectively^84^. These reporter systems are based on an inactive GFP-expressing cassette containing recognition sites for the I-SceI endonuclease. Transfection of I-SceI expressing plasmid induces a DSB within the cassette, and successful repair of the break by NHEJ or HR restores the functional GFP gene. Therefore, the number of GFP-positive cells, counted by flow cytometry, provides a measure of the NHEJ or HR efficiency. Briefly, cells were transfected with I-SceI-T2A-BFP expressing plasmid using Lipofectamine 2000, according to manufacturer’s instructions (Addgene plasmid #45565 was a gift from Andrew Scharenberg^85^). Cells were analyzed 72 h after transfection on a Gallios Flow Cytometer (Beckman Coulter), and DSB repair efficiency was calculated as the percentage of GFP-positive cells within the BFP-positive population (corresponding to transfected cells). Data were normalized to vehicle-treated cells in each experiment.

### Comet assay

Alkaline comet assays were performed using the Comet Assay Kit from R&D Systems (# 4250- 050-K), following manufacturer’s protocol. MIAPaCa-2 and PANC-1 cells were seeded in 6- well plate at a density of 1.25 × 10^5^ cells per well in complete growth medium. After overnight attachment, cells were treated for 24 h or 48 h, harvested, and resuspended in ice-cold PBS 1X. Cells were then mixed with low-melting agarose and immediately loaded on comet slides, followed by incubation at 4°C for 30 min. Slides were then incubated in lysis solution for 1 h and in alkaline unwinding solution for another hour. Alkaline electrophoresis was carried out at 4°C at 19 V for 30 min. Finally, slides were stained with DAPI and imaged using a Thunder microscope (Leica) equipped with a 10× objective lens. Image analysis was performed using OpenComet software within ImageJ^86^.

### Immunoprecipitation of O-GlcNAcylated proteins with RL2-functionalized Dynabeads

Dynabeads M-270 Epoxy (Thermo Fisher Scientific Inc.) were functionalized using the RL2 (ab2739, abcam; 1:1000) antibody, following the protocols implemented by Lagundzin et al^87^. The coupling reaction ratio used was 10 μg of antibody to 0.75 mg of Dynabeads.

For each sample, we immunoprecipitated O-GlcNAcylated proteins by incubating 1 mg of whole protein extract from both untreated and treated MIAPaCa-2 cells. This was done using 0.75 mg of RL2 functionalized Dynabeads M-270 Epoxy (RL2-Dynabeads), following the manufacturer’s instructions. In parallel, a mock-IP was done using only Dynabeads as a negative control. The protein extracts and RL2-Dynabeads were incubated overnight at +4°C under agitation. The following day, beads were first pelleted and then washed with a buffer containing 10 mM Hepes pH 7.4, 10 mM KCl, 50 mM NaCl, 1 mM MgCl_₂_, and 0.05% NP-40, and then with a buffer containing 10 mM Hepes pH 7.4, 10 mM KCl, 0.07% NP-40. Finally, the samples were eluted from the beads by incubation for 10 min with a buffer containing 0.5 M NH_₄_OH and 0.5 mM EDTA, then evaporated using a vacuum evaporator^87^. Each experiment was done in biological triplicate

### In solution trypsin digestion

After denaturation of protein with 8 M urea in 50 mM ammonium bicarbonate (NH_4_HCO_3_), pH 8.8, 10 μg of each sample were reduced and alkylated with 200 mM 1,4-dithiothreitol (DTT) (Sigma-Aldrich), in 50 mM NH_4_HCO_3_ for 1 h at 56 °C, and then with 200 mM 2-iodoacetamide (IAA) (Sigma-Aldrich), in 50 mM NH_4_HCO_3_, for 30 min at RT in darkness. Alkylation reaction was stopped by adding 200 mM DTT in 50 mM ammonium bicarbonate and incubating for 15 min at RT in the dark. The sample was diluted to reduce the urea concentration below 1.4 M and digested using sequencing-grade trypsin (Sigma-Aldrich, Merck Life Science) overnight at 37 °C with a 1:30 (w/w) enzyme-to-protein ratio. Digestion was stopped by the addition of 1% TFA. Whole supernatants were dried down using a vacuum evaporator and stored at −80°C^88^.

### LC-MS/MS analysis

Mass analysis was carried out using Nanoscale liquid chromatography coupled with tandem mass spectrometry. Peptides were separated using an Easy nLC-1000 chromatographic instrument coupled to an Exploris 480 mass spectrometer (ThermoFisher Scientific). Peptides separation was performed using a 15 cm, 75 μm i.d. reversed phase column, in-house packed with 3 μm C18 silica particles (Dr. Maisch). The gradient was generated using mobile phase A (0.1% FA and 2% ACN) and mobile phase B (0.1% FA and 80% ACN). Peptides separation was achieved at a flow rate of 230 nL/min using the following gradient: from 4% B to 44% B in 40 min, from 44% B to 100% B in 8 min, and from 100% B to 0% B in 8 min; then the column was cleaned for 5 min with 100% B. The peptides were subjected to electrospray ionization (ESI) followed by MS/MS. The mass spectrometer operated in DDA mode using a top-6 method. In detail, the MS full scan was 375-1400 m/z with a resolution of 60,000, AGC target ‘’custom’’ and maximum injection time of 50 ms. The mass window for the isolation of the precursor was 1.6 m/z, with a resolution of 30000, an AGC target ‘’custom’’ and a maximum injection time of 120ms; HCD fragmentation was set at normalized collision energy of 30 and dynamic exclusion of 20 s.

Raw files were processed with the MaxQuant software version 2.1.4 using the Andromeda search engine against Human-refprot-isoforms.fasta in the NCBI. Database searches were performed using a precursor mass tolerance of 10 ppm, MS/MS tolerance of 0.02 Da and trypsin cleavage with two missed cleavages allowed. However, carbamidomethylation of cysteine (57.021 Da) was set as a fixed modification, and oxidation of methionine (15.994 Da) have been selected as variable modifications. Potential common contaminants, proteins identified in the decoy database, and proteins only identified by site were excluded. Proteins identified in the Mock-IP were classified as nonspecific interactors and eliminated from the list of RL2 interactors.

Protein quantification was based on the MaxQuant label-free algorithm using unique and razor peptides and the matching-between-run feature selection^41^. Statistical analysis for RL2 enriched proteins was performed using Perseus software version 2.0.11. Data were log_2_ transformed based on LFQ intensity, considering a minimum of 100% valid values in each group.

The values were analyzed using a Student’s t-test to compare proteins enriched in treated cells compared to untreated cells. For statistical analysis and the identification of significant proteins, FDR was set to 0.05%^41^. Protein-protein interaction networks and functional enrichment analyses were performed using IPA Ingenuity software, considering pathways with a p-value < 0.01% as significant.

The MS proteomics data have been deposited to the Proteome Xchange Consortium via the PRoteomics IDEntifications (PRIDE) partner repository with the dataset identifier PXD077403.

### In depth analysis of RUVBL2 glycosylation

500 μg of whole proteins extract of MIAPaCa-2 untreated cells and MIAPaCa-2 cells treated with GEM, isolated as previously reported, were incubated with the antibody against RL2 (ab2739, abcam) at a concentration of 10 μg/mg of proteins, at 4 °C for 4 h on a rotating device. 20 μL of resuspended volume of Protein A/G PLUS-Agarose (Santa Cruz Biotechnology Inc) were added to the mixture and incubated at 4 °C on a rotating device overnight. Beads were washed three times and incubated at 100 °C for 5 min. In parallel, a mock-IP was done using only Protein A/G PLUS-Agarose as a negative control. The resultant immunocomplexes and the relative input were resolved through precast SDS-PAGE (Any kD^TM^ Mini-PROTEAN Precast Protein Gels, Bio-Rad) and visualized using EZBlue Gel Staining Reagent (Sigma-Aldrich, Merck Life Science).

Gel lane referred to immunocomplexes and Mock-IP were sliced according to molecular weight and processed according to the protocol previously reported. Briefly, gel bands were de-stained, in-gel-trypsin digested and purified using SCX tip; resulting peptides were eluted with 0.5% ammonium acetate in 20% acetonitrile and dehydrated in a vacuum evaporator^89^.

We focused our attention on the gel slice corresponding to 50 kDa to confirm that the isoform of RUVBL2 undergoing O-GlcNAcylation is the full-length. This also allowed us to better define the peptide undergoing the modification. The resulting peptides were analyzed according to the protocol outlined in the "LC-MS/MS Analysis" section.

Raw data files were pre-processed with Proteome Discoverer 1.4 (ThermoFisher Scientific). MS/MS data were sought on the Human UniProt database. To estimate the false discovery rate (FDR) of peptide identifications, “Target-decoy PSM validator” node was applied in Proteome Discoverer. Searching parameters were MS error tolerance: 10 ppm; MS/MS error tolerance: 0.02 Da; enzyme specificity: trypsin; maximum number of missed cleavages: 2; taxonomy Human; fixed modifications: Carbamidomethylation (C); variable modification: Oxidation (M), O-GlcNAcylation (HEXNAC) (ST). Only proteins that were identified by two high-scoring tryptic peptides were considered valid. (Xcorr > 2.0 for doubly charged peptides, >2.5 for triply charged peptides, and >3.0 for peptides having a charge state >3 to consider a peptide identification valid)^90^.

### Validation of RUVBL2 O-GlcNAcylation through Western blot

For western blot validation, protein extracts were enriched using Dynabeads functionalized with the antibody anti-RUVBL2 antibody (HPA067966, Sigma-Aldrich, Merck Life Science) following the previously reported protocol. Eluted proteins and the input were evaporated using a vacuum evaporator,resuspended in Laemly buffer, denatured at 100°C for 5 min and resolved by precast SDS-PAGE (Any kD^TM^ Mini-PROTEAN Precast Protein Gels, Bio-Rad). Proteins were electrotransferred by a Transblot turbo system (Bio-Rad) for 30 min according to Bio-Rad protocol. Membrane was incubated with primary antibody RL2 (ab2739, abcam; 1:1000); anti-RUVBL2 antibody (HPA067966, Sigma-Aldrich, Merck Life Science; 1:10000), anti-vinculin (#13901, Cell Signaling Technology Inc.; 1:10000).

The detection of primary anti-RUVBL2 was done by using Clean-Blot™ IP Detection Reagent (HRP) (ThermoFisher Scientific). This reagent is optimized for post-immunoprecipitation Western blot detection of primary antibodies without interference from denatured IP antibody fragments. The detection of primary anti-RL-2 was performed with a horseradish peroxidase-conjugate secondary antibody anti-mouse (sc-7074; Santa Cruz Biotechnology Inc.; 1:2000).

The detection of primary anti-vinculin antibody was performed with a horseradish peroxidase-conjugate secondary antibody anti-rabbit (#7074; Cell Signaling Technology Inc.;1:3000) Immunoblots were developed using the SuperSignal West Femto ECL substrate (Pierce, ThermoFisher Scientific). Corresponding images were acquired by Alliance 2.7 (UVITEC, Eppendorf, Milan, Italy). Densitometric analysis was done by Alliance 1D fully automated software.

### Computational analysis of interaction energy at the monomer-monomer interface in the RUVBL2-RUVBL1 protein hexamer

The stability of the hetero-hexameric wild-type protein complex, consisting of three RUVBL1 alternated to three RUVBL2 subunits (PDB_ID: 9emc.pdb^48^), and that of the corresponding hetero-examer carrying a T81-O-GlcNAcylated residue at the RUVBL2 subunits, were compared by estimating the interaction energy at the interface between the composing monomers for both protein complex models. Specifically, the proteins complexes were solvated through the employment of the CHARMM-GUI ‘Solution Builder’ tool^91–93^, and for the glycosylated complex one N-acetyl-glucosamine molecule was attached through O-GlcNAcylation to the T81 residue of the RUVBL2 subunits (chains D, E and F of the cryo-EM protein complex 9emc.pdb). The resulting solvated models were minimized with a molecular dynamics (MD) simulation employing NAMD2^94–96^.

Concerning the molecular dynamics simulation, the equilibration script for NAMD2 generated by CHARMM-GUI was used: the system was set to operate at a temperature of 303.15K, kept constant by means of a Langevin thermostat, with an integrator time step of 2fs. Starting from this system, 500 conjugate gradient minimization steps followed by 1000 equilibration steps were performed by using NAMD2 with the force field CHARMM36m, according to previously described protocols^92,95–97^.

To verify and correct the presence of clashes and other problems in the generated MD relaxed protein complex models, the FoldX ‘RepairPDB’ tool was employed to solve putative steric problems. The “RepairPDB” tool operates by optimizing side chain atom positions to minimize the overall energy of the protein^98^. Because the FoldX suite does not include carbohydrate parameters, the tool removes the glycosylations from the protein. To remedy this, the protein complex model was re-glycosylated in ChimeraX by superpositioning the pre- and post-repair models and bonding the N-acetylglucosamine (GlcNAc) moieties extracted from the pre-repair model to their respective threonine residues with the ‘Adjust Bonds’ tool in the post-repair model.

At last, the free energy of binding between monomers was evaluated with the Rosetta ‘InterfaceAnalyzer’ tool as previously described^99–102^, specifically between the three RUVBL2 chains D, E and F (glycosylated or not glycosylated at T81) and the three RUVBL1 chains A, B, C (which do not undergo glycosylation).

The -include_sugars and -alternate_3_letter_codes pdb_sugar.codes options were used to enable Rosetta recognition of carbohydrate residues in the model. During model preparation, a limited number of non-essential atoms not supported by the applied parameterization were excluded from subsequent calculations.

Rosetta energy estimations do not employ physical units such as kcal/mol or kJ/mol, and are instead based on special units called REU (Rosetta Energy Units), which are explained at https://www.rosettacommons.org/docs/latest/rosettabasics/Units-in-Rosetta).

### ATP-Mg redocking and docking analyses

Redocking and docking analyses were performed to detect the impact of RUVBL2 T81 O-GlcNAcylation on the affinity of ATP-Mg for the protein: the model protein structure examined (9emc.pdb) was co-crystallised with ATP-Mg, therefore ATP-Mg was redocked in the non-glycosylated model in order to benchmark the behaviour of the docking algorithm (validation by re-docking^103^).

The redocking was performed in AutoDock Tools with AutoDock4^103,104^: the grid was centered on coordinates [158.81, 131.398, 105.236] and consisted of 60, 46 and 46 grid points at 0.275Å spacing. The ligand’s torsional tree was restricted at the purine-ribose bond, in order to deter twists of the purinic ring. The starting position of the molecule was randomised, and 5000000 evaluations were performed on a population of size 150 with 30000. The same protocol was employed in docking ATP-Mg on the glycosylated protein. A visual evaluation of the poses suggested by Autodock was performed in PyMol^103,105,106^.

### Cell Transfection and analysis

For transient expression of wild-type (WT) or T81A mutant RUVBL2 (MUT), we used vectors encoding N-terminal HA-tagged RUVBL2 and an EGFP-puromycin resistance cassette, enabling visualization of transfected cells by green fluorescence and selection with puromycin (pRP[Exp]-EGFP/Puro-CMV>HA/hRUVBL2, #VB240313-1176tud, pRP[Exp]-EGFP/Puro-CMV>HA/hRUVBL2Mut, # VB240313-1175efv, VectorBuilder, GmbH Neu-Isenburg, Germany). In particular, HEK293T cells were cultured in high-glucose Dulbecco’s Modified Eagle Medium (DMEM) supplemented with 2 mM L-glutamine, 100 U/ml penicillin, 100 μg/ml streptomycin, and 10% fetal bovine serum (FBS). Cells were maintained under standard culture conditions at 37 °C in a humidified atmosphere containing 5% CO_₂_. For transient transfection, cells were seeded in 6-well or 96-well plates to achieve 70-80% confluency at the time of transfection. Vectors (Empty Vector (EV), RUVBL2 WT and RUVBL2 MUT) were transfected using the TransIT-X2 Dynamic Delivery System (MIR6004; Mirus Bio, Sigma-Aldrich, Merck Life Science) according to the manufacturer’s protocol. Following transfection, cells were treated with or without 1 μM GEM and subsequently processed for western blot analysis (6- well plates), for cell viability assays using the Incucyte live-cell imaging system or nuclear-to-cytoplasmic HA-RUVBL2 fluorescence analysis (96-well plate).

### Incucyte live cell analysis

HEK293T were seeded on a 96-well plate at 5 × 10^3^ cells per well. After 24 h, cells were transfected with vehicle (NO DNA), EV, RUVBL2 WT and RUVBL2 MUT. Then, plates were transferred to the Incucyte S3 Live-Cell Imaging system (Sartorius) for incubation and analysis of cell proliferation post-transfection over 72 h. After 48 h from seeding, cells were treated with or without 1 μM GEM. Images were captured in brightfield and green channel every 6 h using a 10× objective. Analysis was performed with the basic analyser in the Incucyte software.

### Synergy analysis

Drug interaction between FR054 and olaparib was evaluated in MIAPaCa-2 and PANC-1 cells using a dose-response matrix design. Cells were seeded in 96-well plates and treated with increasing concentrations of FR054 and olaparib, either as single agents or in combination, according to the indicated concentration matrix. After 48 h of treatment, cell growth was assessed by crystal violet staining. Briefly, cells were fixed with 4% paraformaldehyde for 10 min, stained with 0.5% (w/v) crystal violet solution for 1 h, extensively washed with PBS 1X, air-dried, and the bound dye was solubilized in 1% SDS solution. Absorbance was measured at 570 nm using VICTOR X3 multimode plate reader (PerkinElmer). Growth inhibition was calculated relative to CTRL. Dose-response data were analysed using the SynergyFinder platform (version 3.0) based on the Bliss independence model. Synergy landscapes were generated to visualize the interaction between FR054 and olaparib. Bliss synergy scores were interpreted as follows: positive values indicate synergistic interactions, values close to zero indicate additive effects, and negative values indicate antagonistic interactions. The maximum Bliss synergy score for each dose-response matrix was used to quantify the highest degree of synergy achieved by the drug combination.

### Zebrafish Xenograft experiment

For zebrafish xenograft assays, MIAPaCa-2 cells were cultured in complete growth medium and seeded at 7.5 × 10^5^ cells per 100-mm dish. After 24 h, cells were exposed to the indicated treatments for a further 24 h. Treated cells were labeled with CellTracker Green CMFDA (5 μM; C7025, Thermo Fisher Scientific) for 1 h according to the manufacturer’s instructions, harvested, and resuspended in sterile PBS 1X. Cell suspensions were kept on ice until microinjection of approximately 200 cells in 1 nL into the perivitelline space of 48 h post-fertilization (hpf) embryos, which were then maintained at 34 °C. Fluorescence imaging was performed at 2 h post-injection (hpi) to assess initial engraftment and at 48 hpi to evaluate tumor cell persistence and dissemination using stereo microscopes (Leica).

### Flow Cytometry analysis

Following the indicated treatments and γ-irradiation, cells were fixed with Fix Buffer I and incubated at 37 °C for 10 min. An unstained control containing 3 × 10^6^ cells was included. Following fixation, samples were washed twice with PBS 1X and once with PBS 1X containing 0.02% saponin. For fluorescent cell barcoding, cells were resuspended in PBS 1X-0.02% saponin and maintained on ice before the addition of either PB or PO barcoding dye. Single-stained compensation controls were prepared by adding each dye to separate wells containing unstained cells. All samples were incubated for 20 min at RT in the dark, washed twice with flow wash buffer, and barcoded samples were subsequently pooled. Cells were then permeabilized with cold permeabilization buffer and stored at −80 °C. On the following day, samples were thawed and washed with flow wash buffer. Intracellular staining was performed for 30 min at RT, followed by two additional washes. Finally, cells were resuspended acquired using a flow cytometer.

### Single-cell RNA-seq data processing and analysis

The scRNA-seq data from pre- and post-chemotherapy primary and metastatic pancreatic adenocarcinoma tumors^57^ was downloaded from Gene Expression Omnibus (GEO, accession number GSE205013). The data was read into an anndata object (v. 0.11.1)^107^ and went through quality filtering with Scanpy (v. 1.10.3)^108^, retaining cells with < 25 % mitochondrial and < 30,000 total counts and n_genes_by_counts within the 2nd-98th percentile range to exclude cells with unusually low or high number of detected genes. Gene counts were normalized with scanpy.pp.normalize_total with default parameters and log transformed using log(x+1). Highly variable genes were identified (n=3000), and data were scaled to zero mean and unit variance. Principal component analysis was performed using scanpy.tl.pca (n=35 principal components), followed by neighborhood graph construction with scanpy.pp.neighbors (n=75 neighbors). Leiden clustering was initially performed at a resolution of 0.3. The epithelial clusters were subsequently subclustered at a resolution of 0.5. Resulting clusters were annotated based on the expression of cell type-specific marker genes (Supplementary Table 10).

Further processing was done using Scanpy v. 1.9.8. Previously unannotated cell populations were classified as erythrocytes based on the expression of erythrocyte-associated marker genes (HBA1, HBA2, HBB). For the remaining unknown cluster, differential expression analysis against all other cells was performed using scanpy.tl.rank_genes_groups. Differentially expressed genes (Benjamini-Hochberg pAdj < 0.05) were subjected to rank-based gene set enrichment analysis using GSEApy v. 1.1.3^109^. Based on significant enrichment of MSigDB Hallmark^17^ TNF signaling via NF-кB (NES = 2.63) and Inflammatory Response (NES = 2.50) gene sets (pAdj < 0.05, two-sided Wilcoxon rank-sum test), this cluster was annotated as unknown immune cells. Two-dimensional UMAP embeddings were generated using scanpy.tl.umap.

### Gene expression analysis

RNAseq analysis for MIAPaCa-2 and BxPC3 cell lines have been described in ^17^. The RNA-seq data have been deposited at the NCB-GEO database, accession number GSE223303.

### Statistics and reproducibility

Statistical analyses and data visualization were performed using GraphPad Prism v8.0.2 (GraphPad Software Inc., La Jolla, CA, USA), Python statistical libraries (i.e., scipy, pingouin, statsmodels) and Python graphical libraries (i.e., matplotlib). Data are presented as mean ± s.d., mean ± s.e.m., or median with interquartile range, as indicated in the figure legends. P values < 0.05 were considered statistically significant. Statistical analyses were performed using one-way or two-way analysis of variance (ANOVA), followed by Tukey’s or Bonferroni’s multiple-comparisons test when appropriate and specified in the figure legends. For non-parametric comparisons involving multiple groups, the Kruskal-Wallis test followed by Dunn’s multiple-comparisons test was used. Cell-level statistical comparisons of gene set activity between untreated and treated ductal and acinar cells were performed using a two-sided Wilcoxon rank-sum test. The statistical tests applied are specified in the corresponding figure legends. No statistical method was used to predetermine sample size. Sample sizes were based on standard practice in the field and previous studies. No data were excluded from the analyses. Experiments were not randomized, and investigators were not blinded during data acquisition. Imaging-based analyses were performed using standardized acquisition settings and automated analysis pipelines to minimize potential bias.

## Data availability

The publicly available Pancreatic cancer single-cell RNA-sequencing data analyzed in this study have been deposited in the Gene Expression Omnibus (GEO) database under accession code GSE223303 (https://www.ncbi.nlm.nih.gov/geo/query/acc.cgi?acc=GSE223303).

## Contributions

F.C. conceived and designed the study, supervised the overall work, interpreted the data and wrote the manuscript. B.Z. performed most of the experiments, data analysis, figure preparation and review of the manuscript. G.T. performed several experiments and data analysis. S.L. performed the comet assays under the supervision of S.B. C.B. performed the Zebrafish experiments. G.C.M.P. and C.L.P. performed the computational modelling and structural analysis of RUVBL2. K.D. performed the g-irradiation experiments. S.H. and T.S. performed the single-cell transcriptomic re-analysis from public data. A.V. performed the GSVA analysis. A.U. supervised the bioinformatic and transcriptomic analyses performed by the Finnish/Norwegian group. E.D.B. contributed to the initial Operetta-based imaging analyses. M.L.C. and M.P. performed the proteomic analyses and the identification of RUVBL2 O-GlcNAcylation, under the supervision of D.S., who also contributed to proteomic data interpretation and analysis. L.T. and B.LF. synthesized FR054. F.C., S.B., B.LF., C.B., A.U. and D.S. secured funding supporting the study. All authors contributed to data interpretation, discussed the results and reviewed the manuscript.

## Acknowledgements

This work was supported by the University of Milano-Bicocca through FAR grants 2026-ATE- 0054, 2025-ATE-0426, 2024-ATE-0434, 2023-ATE-0359 and 2021-ATEQC-0006 to F.C. and 2025-ATE-0193 to B.LF. F.C. and D.S. acknowledge support from the Italian Ministry of University and Research (MUR), 2022ZBZFX3 PRIN-2022. B.Z. and G.T. were supported by PhD fellowships funded by MUR. S.B. acknowledges support from the Italian Ministry of University and Research (MUR), under the 20222KSN2N PRIN-2022 grant. A.U. is a K. Albin Johansson Cancer Research Fellow and a Tampere Institute of Advanced Study Research Fellow. A.U. acknowledges support from the Finnish Cancer Institute, the Research Council of Finland (project no. 349314), Cancer Foundation Finland, and the Norwegian Cancer Society (project nos. 198016-2018 and 273672-2023). K.D. acknowledges support from the Norwegian Cancer Society (project no. 273672-2023). T.S. acknowledges support from Cancer Foundation Finland and the Research Council of Finland (project no. 349314).

---

**Supplementary Figure 1.** Cyclin A and γH2AX subpopulation analysis in MIAPaCa-2 and PANC-1 cells following FR054, GEM, and COMBO treatment. a, b, Quantification of Cyclin A/γH2AX subpopulations in MIAPaCa-2 (a) and PANC-1 (b) cells. Bar plots show the percentage of cells (mean ± s.d., *n*=3 independent biological replicates) distributed across four subpopulations (Cyclin A-low/γH2AX-low, Cyclin A-high/γH2AX-low, Cyclin A-low/γH2AX- high and Cyclin A-high/γH2AX-high) following treatment with vehicle (CTRL), 500 μM FR054, 1 μM gemcitabine (GEM), or their combination (COMBO) at 24 h and 48 h post- treatment. Statistical significance in a, b was determined using two-way ANOVA followed by Tukey’s multiple-comparisons test. Exact P values are indicated in the plots; ns, not significant.

**Supplementary Figure 2**. **Time-dependent modulation of** γ**H2AX following ionizing radiation and FR054 treatments. a**, Flow-cytometry histograms showing the γH2AX fluorescence intensity in MIAPaCa-2 cells exposed to increasing radiation doses (1, 3, or 5 Gy) and collected at the indicated time points (0 h, 1 h, 2 h, 24 h, 48 h, and 72 h post-irradiation). **b**, Dose- dependent γH2AX level after 48 h FR054 treatment in the indicated cells. Flow-cytometry histograms showing the distribution of the γH2AX fluorescence intensity in cells treated with increasing concentrations of FR054 (25-300 μM) compared to untreated control (CTRL).

Supplementary Table 1. Differentially expressed genes in MIAPaCa-2 cells following 500 μM FR054 (MF), 1 μM GEM (MG) or combined treatment (MFG) (1488). Genes significantly differentially expressed after 48 h of treatments, relative to untreated control cells (log_₂_FC| ≥ 2; FDR ≤ 1 × 10^-5^).

Supplementary Table 2. Differentially expressed genes in BxPC3 cells following 500 μM FR054 (BF), 1 μM GEM (BG) or combined treatment (BFG) (1026). Genes significantly differentially expressed after 48 h of treatments, relative to untreated control cells; log_₂_FC| ≥ 2; FDR ≤ 1 × 10^-5^).

Supplementary Table 3. Differentially expressed genes (DEGs) shared between MIAPaCa-2 and BxPC3 cells following 500 μM FR054 (MF, BF respectively), 1 μM GEM (MG, BG respectively) or combined treatment (MFG, BFG respectively); log_₂_FC| ≥ 2; FDR ≤ 1 × 10^-5^.

Supplementary Table 4. List of the 275 DEGs shared in combined treatment of 500 μM FR054 and 1 μM GEM between MIAPaCa-2 (MFG) and BxPC3 (BFG) cells. List of 275 differentially expressed genes compared between MIAPaCa-2 and BxPC3 cells, including the log_₂_ fold- change values in each cell line and their classification as concordantly regulated (270 genes) or oppositely regulated (5 genes).

Supplementary Table 5. DNA damage-related differentially expressed genes in MIAPaCa-2 cells following 500 μM FR054 (MF), 1 μM GEM (MG) or combined treatment (MFG) (320 genes). List of 320 differentially expressed genes associated with DNA damage-related C2 gene sets in MIAPaCa-2 cells treated for 48 h with FR054, GEM or their combination, relative to untreated control cells (CTRL). Values indicate log_2_ fold changes. Blank cells indicate genes that did not meet the differential-expression criteria in the corresponding comparison (|log_₂_FC| > 2; FDR < 0.05).

Supplementary Table 6. List of the identified proteins from the immunoprecipitation of untreated and treated MIAPaCa-2 cells using the RL2 antibody, MaxQuant output results. The table details for each identified protein: protein ID, protein name, gene name, LFQ intensity for each biological replicate, number of peptides, number of razor unique peptide, number of unique peptides, sequence coverage [%], razor peptide sequence coverage [%], unique sequence coverage [%], mol. weight [kDa], Protein identified only by site, in the reverse decoy and as potential contaminants were excluded from the analysis.

Supplementary Table 7. Detailed information on the differentially enriched proteins in treated cells compared to untreated cells, Perseus output results. The table details for each identified protein: protein ID, protein name, gene name, number of peptides, number of razor unique peptides, number of unique peptides, sequence coverage [%], razor peptides sequence coverage [%], unique sequence coverage [%], mol. weight [kDa], and LFQ intensity for each replicate of analyzed sample, Significant proteins (FDR<0,05) are marked with*, Difference of treated cells vs untreated and log(p-value) of the difference.

Supplementary Table 8. Computational analysis of RUVBL1-RUVBL2 complex stability and ATP-Mg binding. Rosetta InterfaceAnalyzer metrics and AutoDock-predicted ATP-Mg binding affinities for the non-glycosylated and Thr81-O-GlcNAcylated RUVBL1-RUVBL2 complexes.

Supplementary Table 9. Cell subtype annotation marker genes. Canonical marker genes used to identify immune, stromal and epithelial cell populations, including pancreatic endocrine, acinar and ductal cells, in the analysed single-cell transcriptomic dataset.

Supplementary Table 10. Hallmark DNA repair geneset. List of the 150 genes included in the Hallmark DNA Repair gene set used for gene set enrichment and variation analyses.

**Extended Data Figure 1.**
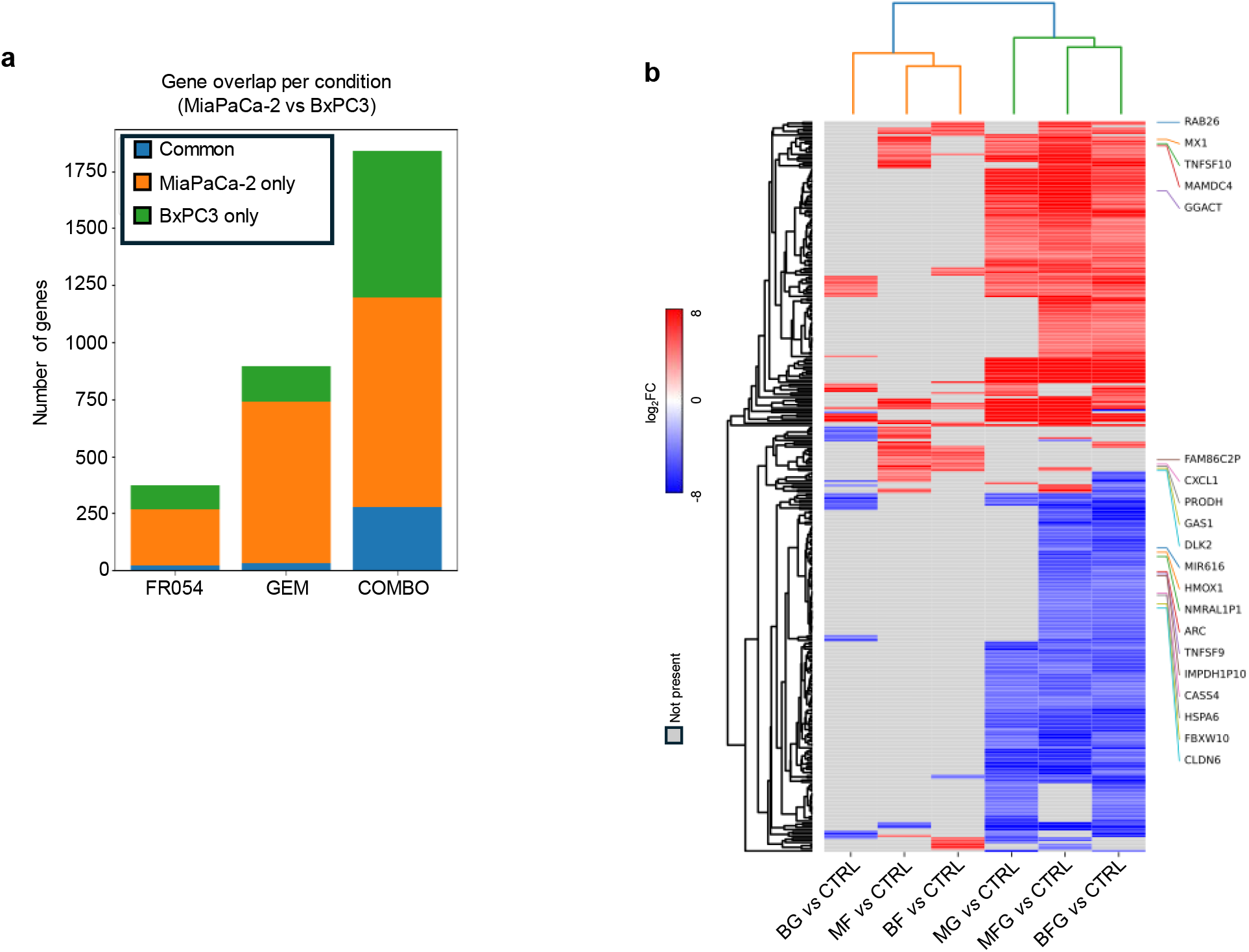
Comparison of treatment-induced transcriptional changes in MIAPaCa-2 and BxPC3 pancreatic cancer cells. **a**, Number of differentially expressed genes (DEGs) shared between MIAPaCa-2 and BxPC3 cells or detected exclusively in either cell line following treatment for 48 h with 500 μM FR054 (MF, BF), 1 μM gemcitabine (MG, BG), or their combination (MFG, BFG), relative to the corresponding vehicle-treated control (CTRL). DEGs were defined by an absolute log_2_ fold change (log_2_FC) > 2 and a false discovery rate (FDR) < 0.05. Bars indicate genes common to both cell lines (dark grey), specific to MIAPaCa-2 cells (red) or specific to BxPC3 cells (blue). **b**, Heatmap showing the log_2_FC values of treatment-responsive genes across the six pairwise comparisons: FR054-treated MIAPaCa-2 versus CTRL (MF versus CTRL), FR054-treated BxPC3 versus CTRL (BF versus CTRL), GEM- treated MIAPaCa-2 versus CTRL (MG versus CTRL), GEM-treated BxPC3 versus CTRL (BG versus CTRL), COMBO-treated MIAPaCa-2 versus CTRL (MFG versus CTRL) and COMBO-treated BxPC3 versus CTRL (BFG versus CTRL). Genes are grouped according to their treatment-dependent regulation and overlap between the two cell lines. Red and blue indicate increased and decreased expression, respectively, whereas grey indicates that the gene was not significantly differentially expressed or was not detected in the corresponding comparison. Statistical analysis and DEG selection were performed using the same criteria as in Fig. 1.

**Extended Data Figure 2.**
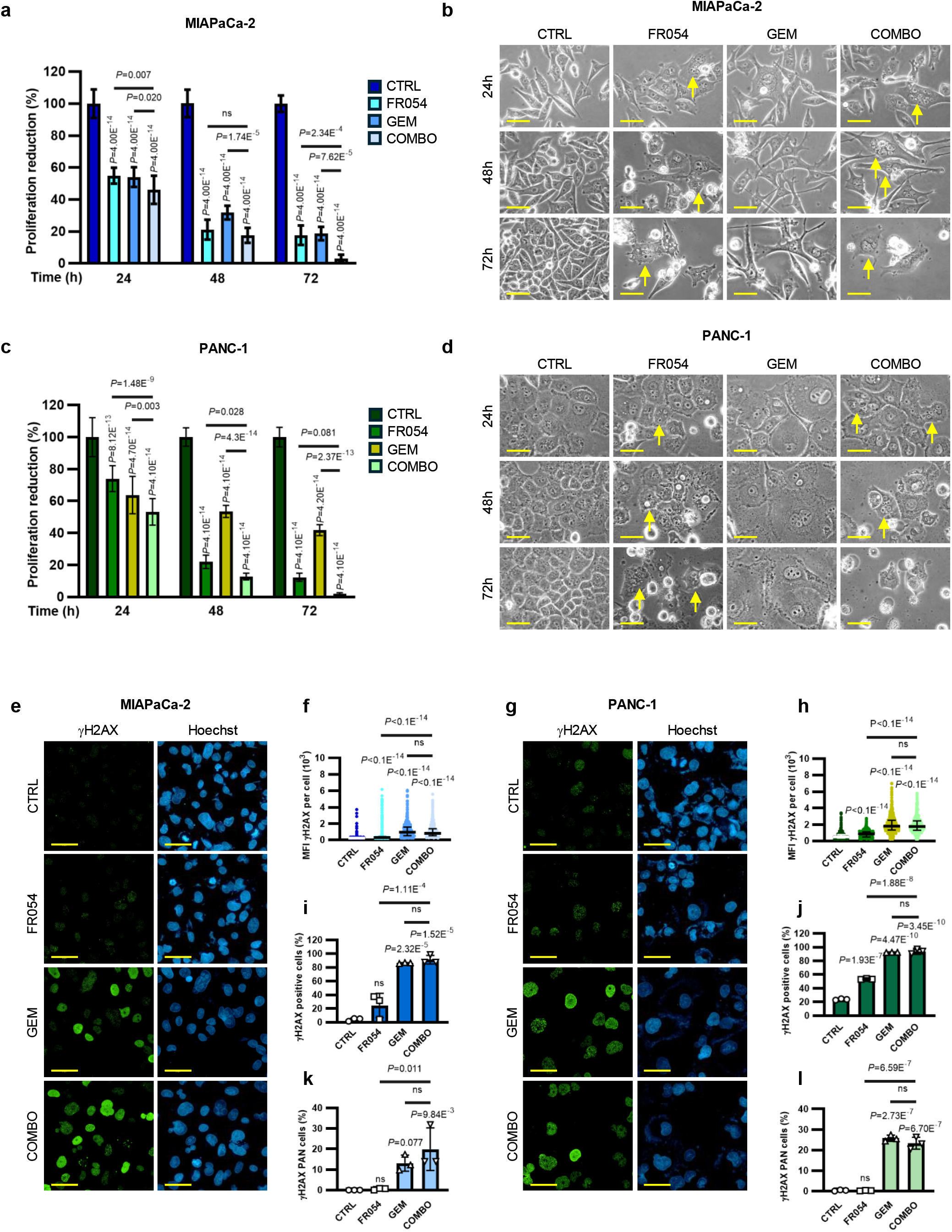
FR054 inhibits cell proliferation, induces multinucleation and increases γH2AX accumulation in PDAC cells. **a, c**, Viable cell number determined by Trypan Blue exclusion in MIAPaCa-2 (**a**) and PANC-1 (**c**) cells treated with vehicle (CTRL), 500 μM FR054, 1 μM gemcitabine (GEM), or their combination (COMBO) for 24 h, 48 h or 72 h. Values are expressed as a percentage of the corresponding time-matched CTRL. The data represent *n*=7–12 (**a**) and *n*=5–13 (**c**) biological replicates. **b**, **d**, Representative phase-contrast images of MIAPaCa-2 (**b**) and PANC-1 (**d**) cells under the conditions described in **a**, **c**. Yellow arrows indicate multinucleated cells. Scale bar, 50 μm (20× magnification). **e**, **g**, Representative immunofluorescence images of γH2AX in MIAPaCa-2 (**e**) and PANC-1 (**g**) cells treated with CTRL, FR054, GEM or COMBO for 24 h. γH2AX is shown in green and nuclei were counterstained with Hoechst (blue). Images are representative of three independent biological replicates (*n*=3) with similar results; scale bar, 50 μm (63× magnification). **f**, **h**, Single-cell quantification of γH2AX mean fluorescence intensity (MFI) in MIAPaCa-2 (**f**) and PANC-1 (**h**) cells using Harmony software after 48 h of treatments as described in **a**, **c**. Scatter dot plots showing individual values, with the median indicated by the central line and the interquartile range (IQR). The number of analysed cells per condition is indicated by the range (MIAPaCa-2: 5075–7951 (**f**); PANC-1: 1593–2597 (**h**)) from three independent biological replicates (*n*=3). **i**, **j**, Percentage of γH2AX-positive MIAPaCa-2 (**i**) and PANC-1 (**j**) cells after 24 h of treatment as described in **a**, **c** (*n*=3 biological replicates). **k**, **l**, Percentage of MIAPaCa-2 (**k**) and PANC-1 (**l**) cells displaying PAN-nuclear γH2AX staining after 24 h of FR054, GEM and COMBO treatment (*n*=3 biological replicates). Data in **a**, **c**, **i**-**l** are presented as mean ± s.d; individual symbols in **i**–**l** represent independent experiments. Statistical significance was determined using two-way ANOVA with Tukey’s multiple comparisons test (**a** and **c**), Kruskal-Wallis test with Dunn’s multiple comparisons test (**f** and **h**) or one-way ANOVA with Tukey’s multiple comparisons test (**i**–**l**). Exact *P* values are reported in the plots; ns, not significant.

**Extended Data Figure 3.**
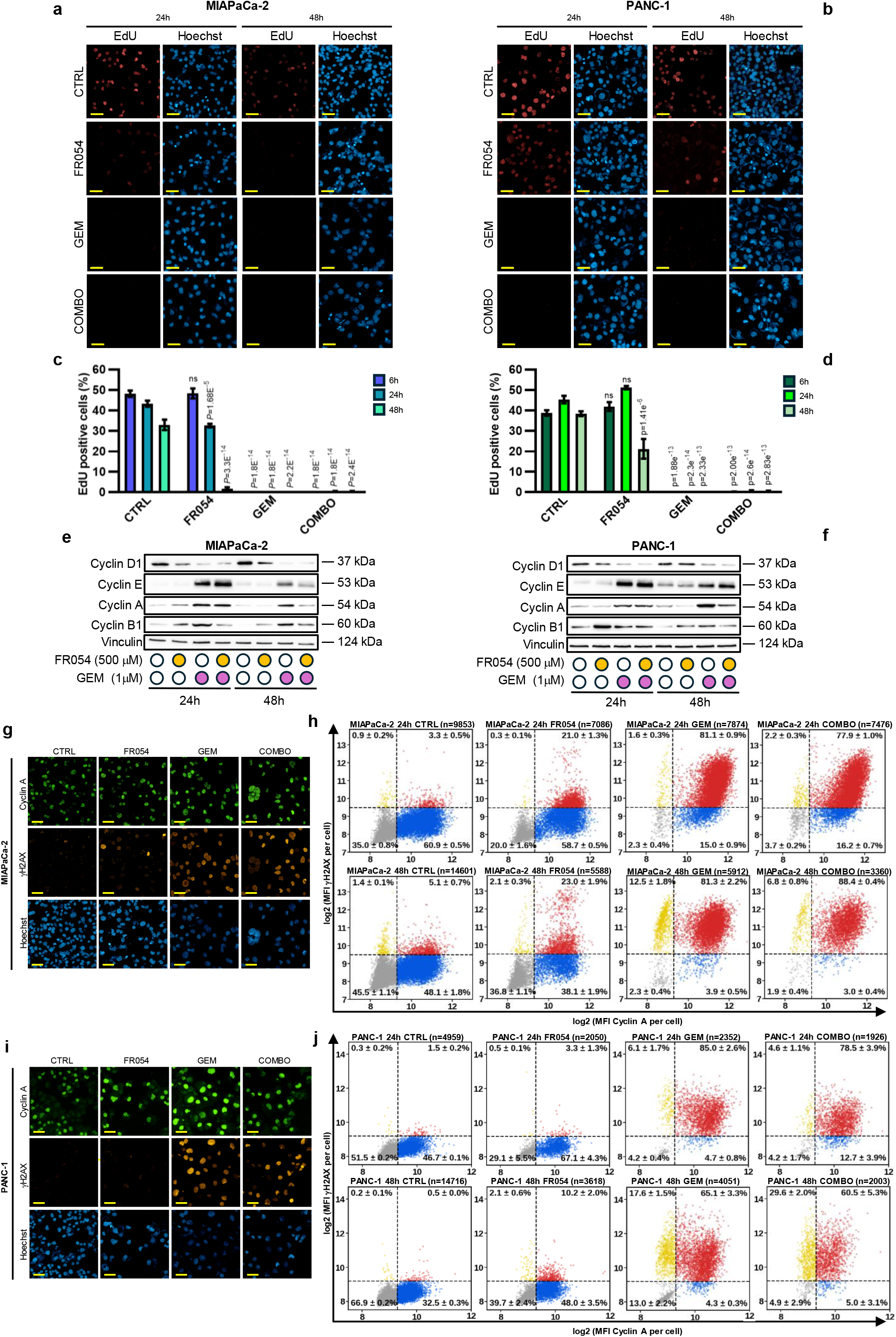
FR054 reduces DNA synthesis and remodels cell-cycle protein expression and the Cyclin A/γH2AX relationship in PDAC cells. **a**, **b**, Representative immunofluorescence images of EdU incorporation in MIAPaCa-2 (**a**) and PANC-1 (**b**) cells treated with vehicle (CTRL), 500 μM FR054, 1 μM gemcitabine (GEM), or their combination (COMBO) for 24 h or 48 h. EdU is shown in red and nuclei were counterstained with Hoechst (blue). Images are representative of three independent biological experiments (*n*=3). Scale bar, 50 μm (40× magnification). **c**, **d**, Percentage of EdU-positive MIAPaCa-2 (**c**) and PANC-1 (**d**) cells after 6 h, 24 h and 48 h of treatment (*n*=3 independent biological experiments). **e**, **f**, Representative immunoblots showing treatment-induced changes in Cyclin D1, Cyclin E, Cyclin A and Cyclin B1 protein levels in MIAPaCa-2 (**e**) and PANC-1 (**f**) cells treated as described in **a**, **b**. Immunoblots are representative of three independent biological replicates with similar results; vinculin was used as a loading control. **g**, **i**, Representative immunofluorescence images showing Cyclin A and γH2AX in MIAPaCa-2 (**g**) and PANC-1 (**i**) cells under the indicated treatment conditions. Cyclin A is shown in green, γH2AX in red and nuclei were counterstained with Hoechst (blue). Images are representative of three independent biological replicates (*n*=3) with similar results. **h, j**, Two-dimensional single-cell density plots showing the relationship between Cyclin A and γH2AX fluorescence intensities in MIAPaCa-2 (**h**) and PANC-1 (**j**) cells after 24 h or 48 h of treatment. Cyclin A and γH2AX mean fluorescence intensities (MFI) were log_2_-transformed. The number of cells analysed in each condition is indicated above the corresponding plot. Data were obtained from three independent biological experiments (*n*=3). Quantification of the identified cell populations and the corresponding statistical analysis are provided in Supplementary Fig. 2. Results are shown as mean ± s.e.m. (**c**, **d**) or mean ± s.d. (**h**, **j**). Statistical significance was determined using two-way ANOVA followed by Tukey’s multiple-comparisons test (**c**, **d**, **h** and **j**). Exact *P* values are indicated in the plots; ns, not significant.

**Extended Data Figure 5.**
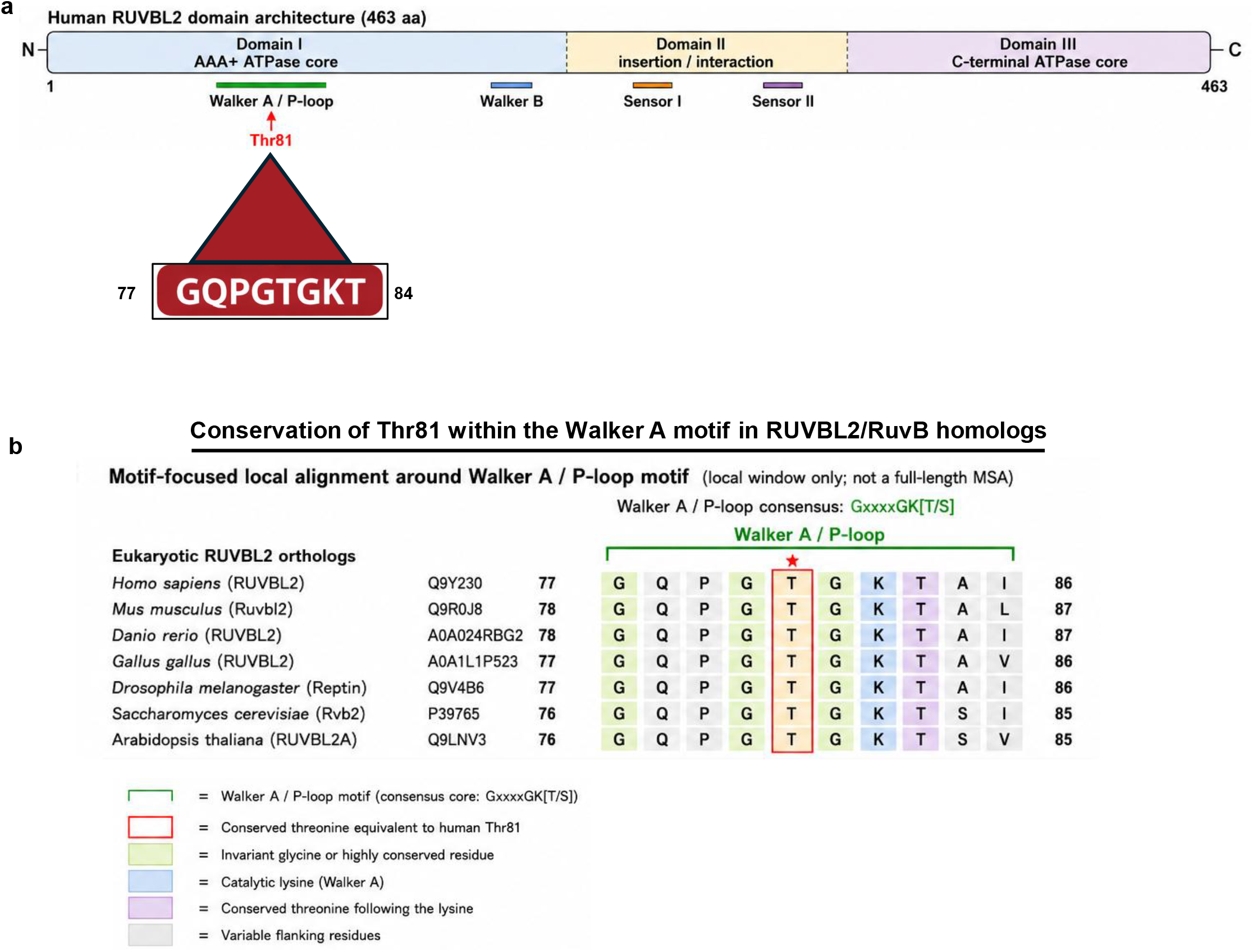
Thr81 is selectively conserved within the Walker A/P-loop region of eukaryotic RUVBL2 orthologues. **a,** Schematic representation of the domain architecture of human RUVBL2 (463 amino acids), comprising domain I, which contains the N-terminal AAA+ ATPase core; domain II, which forms the insertion and interaction domain; and domain III, which contains the C-terminal ATPase core. The positions of the Walker A/P-loop, Walker B, Sensor I and Sensor II motifs are indicated. Thr81 is highlighted within the Walker A/P-loop sequence spanning residues 77-84. **b,** Motif-focused local sequence alignment of the Walker A/P-loop region in representative eukaryotic RUVBL2 orthologues. The alignment represents the local motif only and not a full-length multiple-sequence alignment. The residue corresponding to human RUVBL2 Thr81 is highlighted by a red box. Thr81 is conserved in the analysed eukaryotic RUVBL2 orthologues. The catalytic lysine and the threonine immediately following it are conserved across all analysed sequences. UniProt accession numbers and residue positions are indicated for each sequence.

**Extended Data Figure 7.**
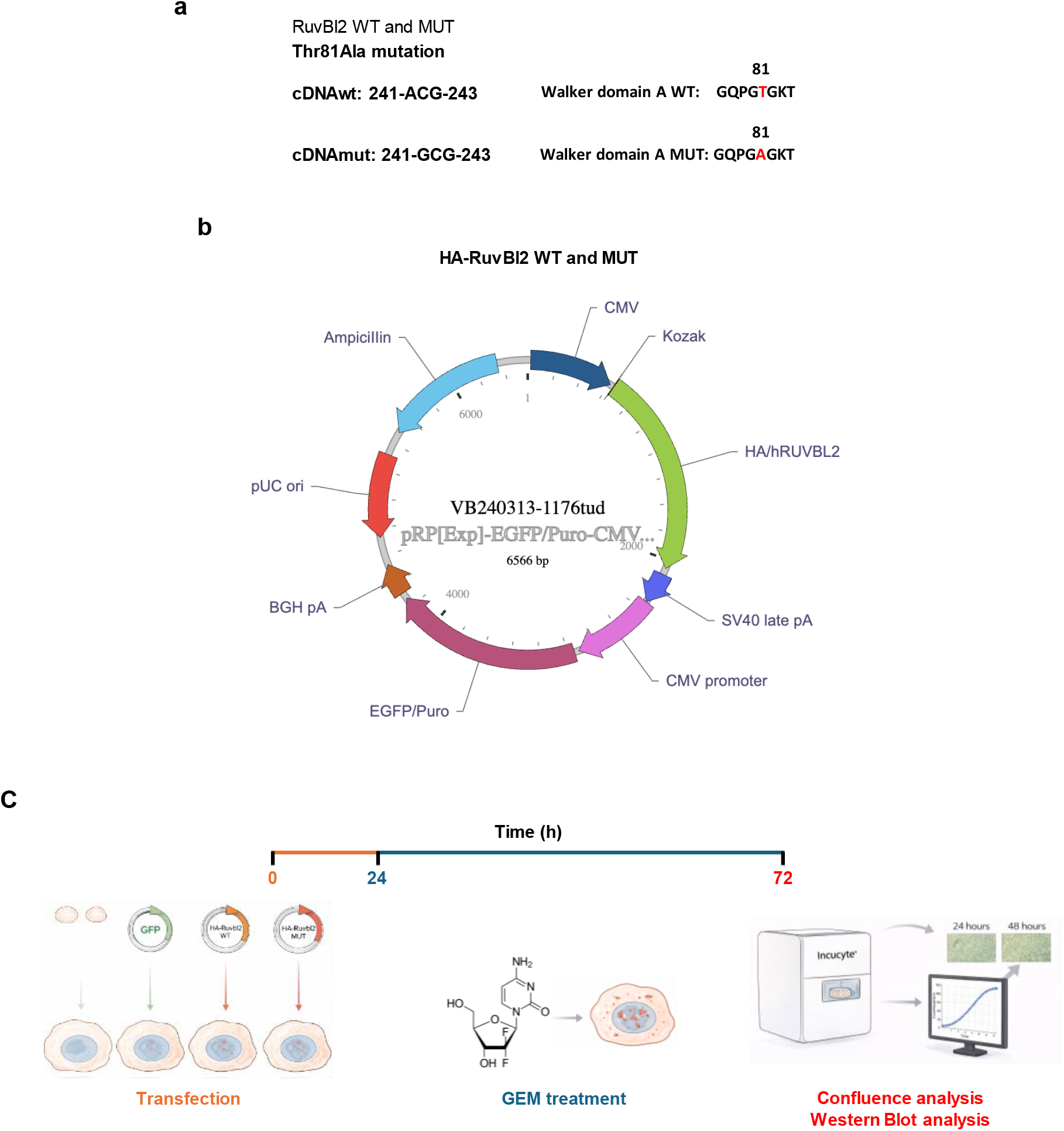
Experimental generation of the RUVBL2 T81A mutant. **a**, The T81A substitution was generated by replacing the wild-type ACG codon at cDNA positions 241-243 with GCG, changing the Walker A sequence from GQPGTGKT to GQPGAGKT. **b**, Schematic representation of the pRP[Exp]-EGFP/Puro-CMV vector used to express HA-tagged wild-type RUVBL2 or the T81A mutant. **c**, Experimental workflow for analysis of cells expressing wild-type or T81A-mutant RUVBL2. Cells were transfected at 0h, treated with gemcitabine (GEM) 24 h after transfection and analysed at 72h by immunoblotting and confluence measurements.

**Extended Data Figure 8.**
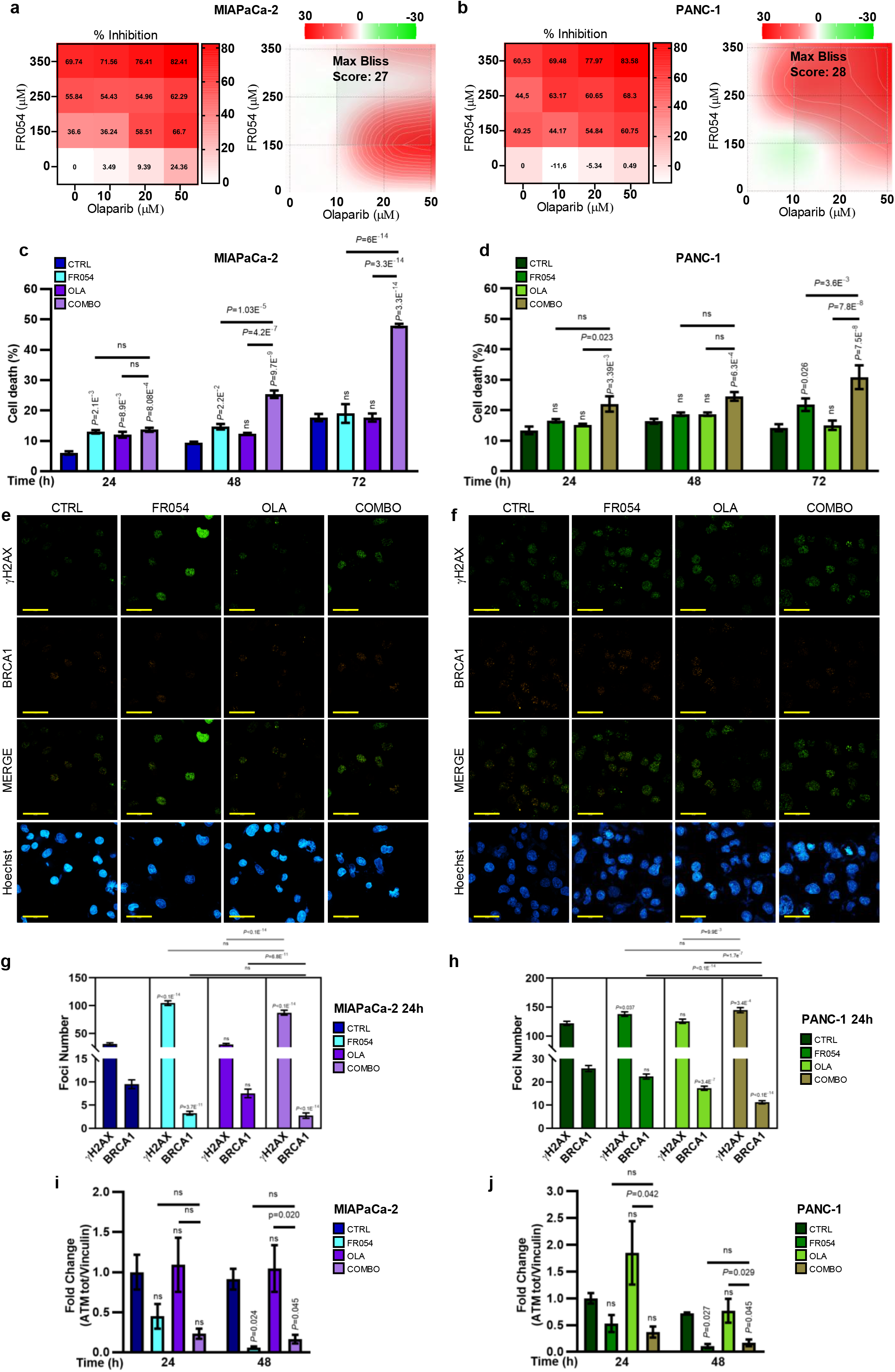
FR054 cooperates with olaparib to increase DNA damage and cell death in PDAC cells. **a**, **b**, SynergyFinder analysis of FR054 and olaparib (OLA) combination in MIAPaCa-2 (**a**) and PANC-1 (**b**) cells. Dose-response matrices show the percentage of growth inhibition induced by the indicated drug concentrations; each matrix value represents the mean of three independent biological replicates. The corresponding Bliss synergy landscapes show positive, additive and antagonistic interactions in red, white and green, respectively. Maximum Bliss synergy scores at 48 h were 27 in MIAPaCa-2 cells (**a**) and 28 in PANC-1 cells (**b**). **c**, **d**, Percentage of dead MIAPaCa-2 (**c**) and PANC-1 (**d**) cells following treatment with vehicle (CTRL), 350 μM FR054, 20 μM olaparib (OLA), or their combination (COMBO) for 24 h, 48 h or 72 h. Data were obtained from *n*=3 (**c**) and *n*≥5 (**d**) independent biological experiments. **e**, **f**, Representative immunofluorescence images of γH2AX and BRCA1 in MIAPaCa-2 (**e**) and PANC-1 (**f**) cells treated with CTRL, FR054, OLA or COMBO for 24 h. γH2AX is shown in green, BRCA1 in red and nuclei were counterstained with Hoechst (blue). Images are representative of three independent biological experiments (n = 3); scale bar, 50 μm (63× magnification). **g**, **h**, Quantification of γH2AX and BRCA1 nuclear foci in MIAPaCa-2 (**g**) and PANC-1 (**h**) cells under the conditions shown in **c**, **d** and **e**, **f**. Bars represent the mean values obtained from *n*=182–243 (**g**) or *n*=362–468 (**h**) nuclei per condition, pooled from three independent biological experiments. **i**, **j**, Densitometric quantification of ATM protein abundance in MIAPaCa-2 (**j**) and PANC-1 (**j**) cells after 24 h or 48 h of treatment as previously described. ATM signals were normalized to vinculin and expressed as fold change relative to the corresponding time-matched CTRL. Quantification was performed from three independent biological experiments (*n*=3). Data are presented as mean ± s.e.m. (**c**, **d** and **g**–**j**). Statistical significance was determined using two-way ANOVA with Tukey’s multiple comparisons test (**c** and **d**), Kruskal-Wallis test followed by Dunn’s post-hoc test (**g** and **h**) or one-way ANOVA with Tukey’s multiple comparisons test (**i** and **j**). Exact *P* values are indicated in the plots; ns, not significant.

