## Supplementary Fig 1 for "PGM3 inhibition rewires RUVBL2-dependent DNA repair and induces a BRCAness-like state in pancreatic cancer cells"

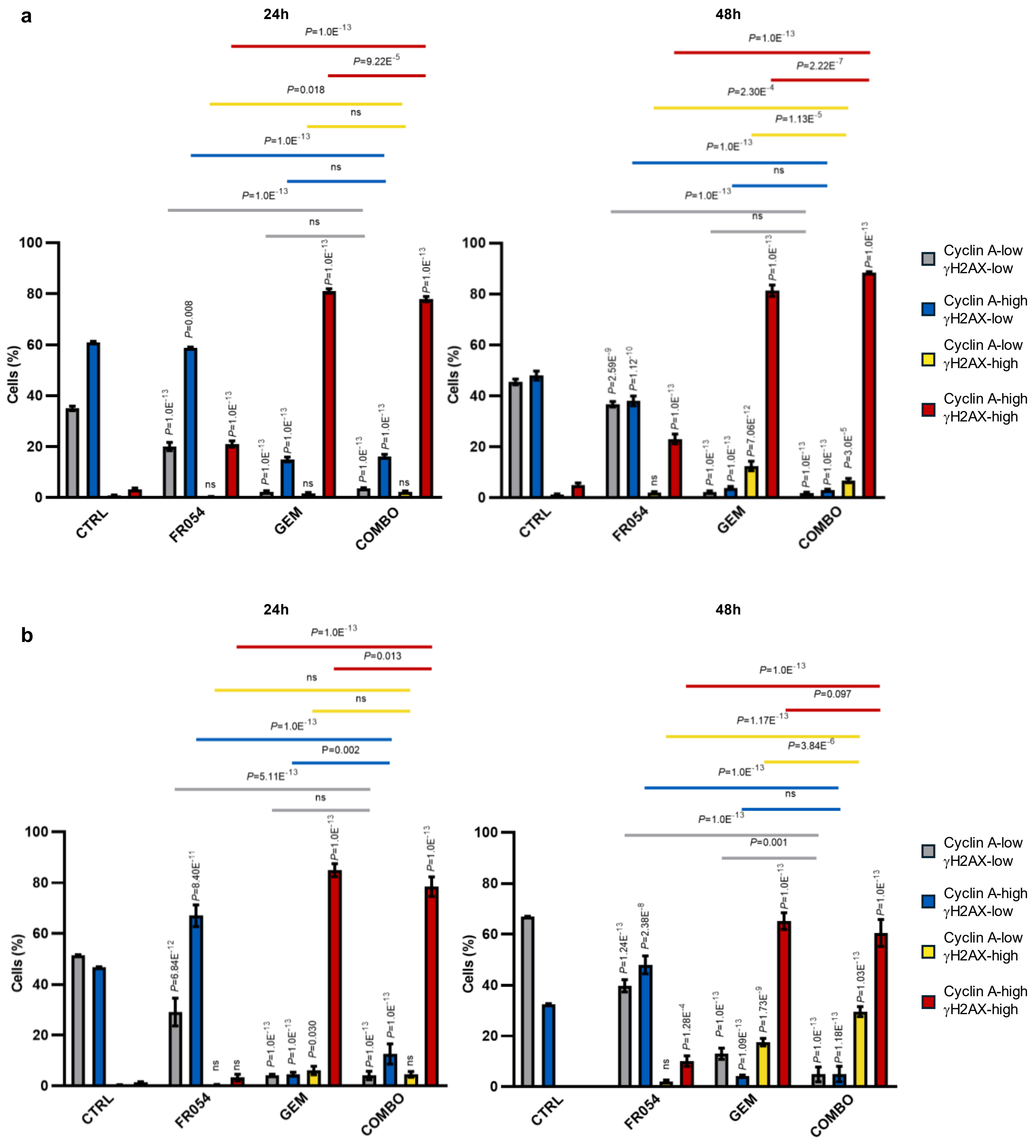


**Supplementary Figure 2. Cyclin A and γH2AX subpopulation analysis in MIA PaCa-2 and PANC-1 cells following FR054, GEM, and combination treatment. a, b,** Quantification of Cyclin A/γH2AX subpopulations in MIAPaCa-2 (**a**) and PANC-1 (**b**) cells. Bar plots show the percentage of cells (mean ± s.d., *n*=3 independent biological replicates) distributed across four subpopulations (Cyclin A-low/γH2AX-low, Cyclin A-high/γH2AX-low, Cyclin A-low/γH2AX-high and Cyclin A-high/γH2AX-high ) following treatment with vehicle (CTRL), 500 μM FR054, 1 μM gemcitabine (GEM), or their combination (COMBO) at 24h and 48h post-treatment. Statistical significance in **a**, **b** was determined using two-way ANOVA followed by Tukey's multiple-comparisons test. Exact P values are indicated in the plots; ns, not significant.
