## Supplementary Fig 2 for "PGM3 inhibition rewires RUVBL2-dependent DNA repair and induces a BRCAness-like state in pancreatic cancer cells"

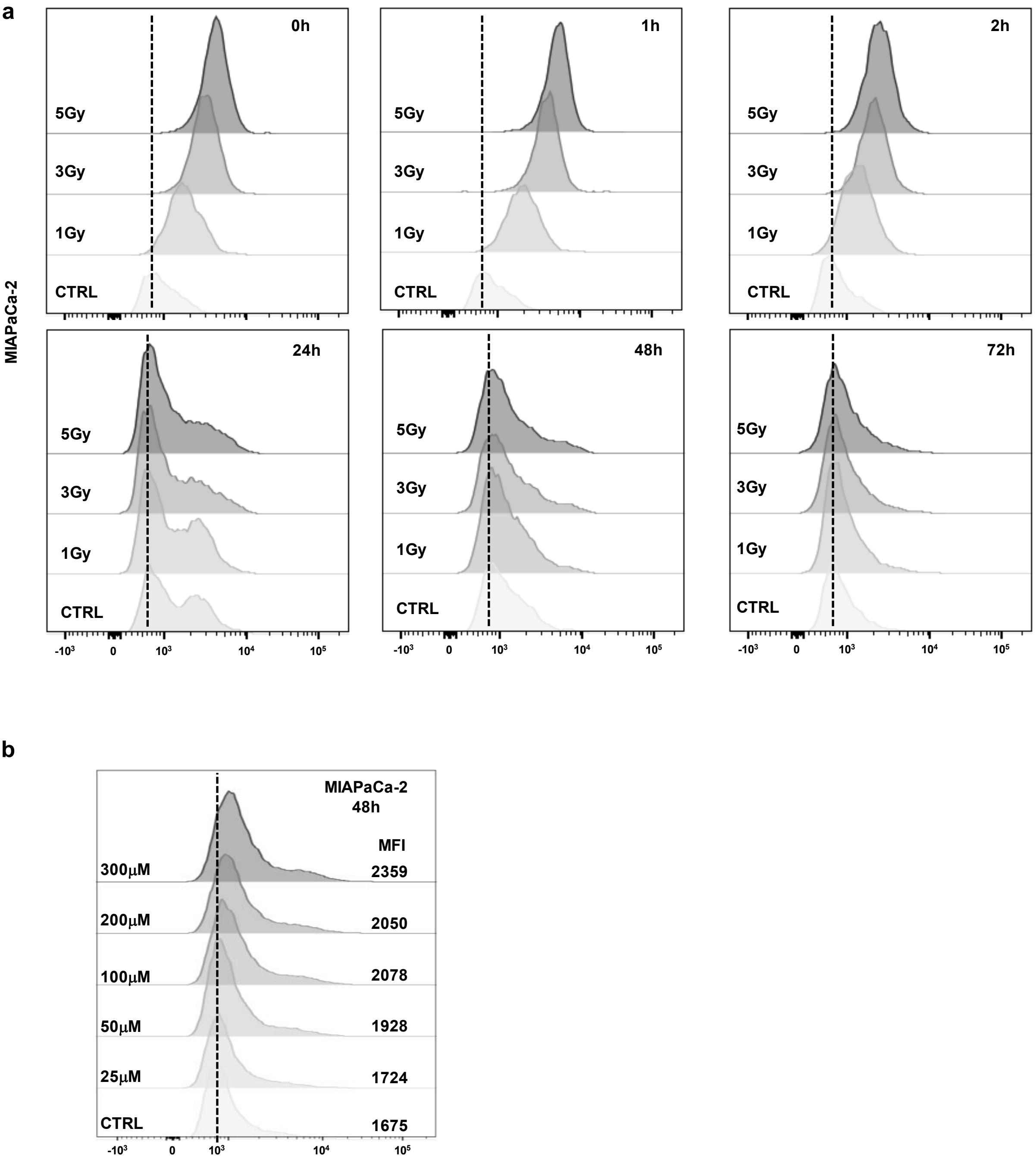


**Supplementary Figure 2** Time-dependent modulation of γH2AX following ionizing radiation. **a.** Flow-cytometry histograms showing the γH2AX fluorescence intensity in MIAPaCa-2 cells exposed to increasing radiation doses (1, 3, or 5 Gy) and collected at the indicated time points (0 h, 1 h, 2 h, 24 h, 48 h, and 72 h post-irradiation). **b**. Dose-dependent γH2AX level after 48 h FR054 treatment in the indicated cells. Flow-cytometry histograms showing the distribution of the γH2AX fluorescence intensity in cells treated with increasing concentrations of FR054 (25-300 μM) compared to untreated control (CTRL).
